# Microscopic collisions govern macroscopic aerotactic spreading through emergent transport coefficients

**DOI:** 10.64898/2026.10.08.757612

**Authors:** N. Ghazi, S. Hirose, K. Funamoto, V. Calvez, C. Anjard, O. Cochet-Escartin, M. Demircigil, J.-P. Rieu

**Affiliations:** Institut Lumière Matière, UMR5306, Université Lyon 1-CNRS, Université de Lyon, 69622 Villeurbanne, France; Graduate School of Biomedical Engineering, Tohoku University, 6-6-12 Aramaki-aza Aoba, Aoba-ku, Sendai, Miyagi 980-8579, Japan; Institute of Fluid Science, Tohoku University, 2-1-1 Katahira, Aoba-ku, Sendai, Miyagi 980-8577, Japan; Laboratoire de Mathématiques de Bretagne Atlantique, UMR 6205, CNRS, Univ Brest, France; Department of Mathematics, University of Arizona, 85721, Tucson, AZ, USA

## Abstract

Collective migration is often attributed to complex cell-cell communication, yet coordinated multicellular dynamics can also emerge in organisms lacking social interactions. Continuum descriptions successfully capture such behaviors but generally rely on phenomenological transport coefficients whose microscopic origin remains elusive. Here we show that these coefficients emerge from cell-cell collisions and can be independently determined from experiments. We investigate the aerotactic dispersal of confined colonies of the solitary *amoeba Acanthamoeba castellanii*, in which oxygen consumption continuously reshapes the local environment to generate self-organized guidance cues. Combining quantitative videomicroscopy, oxygen mapping with phosphorescent sensing films, and oxygen-controlled microfluidics, we directly measure both the oxygen landscape and the aerotactic response governing colony expansion. We then develop a Patlak-Keller-Segel (PKS) -type drift-diffusion model whose transport coefficients are entirely determined from independent experiments and microscopic-to-macroscopic modeling approach, without adjustable parameters. The model quantitatively reproduces the observed dynamics, including the coexistence of a fast ballistic front and a trailing transient accumulation band. We show that both diffusion and aerotactic transport are modulated by the same collision-controlled mechanism through density-dependent changes in cell persistence. More generally, our results establish transport coefficients as measurable emergent properties bridging microscopic cell dynamics and macroscopic collective behavior, providing a predictive framework for oxygen-guided migration in living active matter.

## Introduction

Collective cell migration is a hallmark of living systems, underlying embryonic development, tissue morphogenesis and cancer invasion (1), immune responses (2) or wound healing (3). Collective cell migration and cell coordination is also crucial for microbial colonization (4), and for the foraging strategies used by predators to explore their landscape (5). Although often described as coordinated behavior, collective migration can emerge from remarkably diverse microscopic interactions. In minimal interaction regimes, coordination arises either through direct cell-cell contacts or through self-generated chemical gradients. Several contact-mediated mechanisms have been identified, including Contact Inhibition of Locomotion (CIL) (6), Contact Enhancement of Locomotion (CEL) (7), Contact Following (CF) (8), and Contact Attraction (CA) (9), which, alone or in combination, generate a broad spectrum of collective behaviors (10). At the continuum level, however, these diverse microscopic interaction mechanisms do not necessarily require explicit modeling. Instead, their collective effects can often be coarse-grained into effective transport coefficients that summarize how local interactions shape random motility and directional migration (11–14).

While cells can migrate randomly at short scales, efficient migration in vivo up to large physiological scales is usually guided by directional cues such as diffusible cues called chemoattractants that are generally detected by receptors on the cell surface. The direction and steepness of the chemical gradients trigger the cell migration directed towards or away from the source of the chemoattractant (15). Different organisms employ different strategies to sense such gradients. In swimming bacteria, chemotaxis is classically based on temporal sensing: because concentration differences across the small bacterial cell body are generally weak, cells detect changes in concentration as they move along their trajectories (16). Larger eukaryotic cells, by contrast, are able to directly compare receptor occupancy across the cell body and can therefore perform spatial gradient sensing (17). These strategies are not mutually exclusive, however: eukaryotic cells can also respond to temporal variations in chemical signals (18), and spatial sensing has recently been demonstrated in some bacterial systems (19).

Although externally imposed gradients efficiently guide migration over short distances, they are difficult to maintain over the millimeter-to-centimeter scales encountered during tissue morphogenesis, wound healing or microbial colonization. Self-generated gradients provide an elegant solution to this physical limitation because the migrating population continuously regenerates the guidance cues. The aggregation of starved *Dictyostelium discoideum* (*Dd*) amoebae by periodic waves of cAMP secreted and relayed from cell to cell is an extremely sophisticated mechanism to propagate chemical gradients at distances (20, 21). Another very generic mechanism is the degradation or uptake of an essential molecule by cells which, locally, create a sink. If the cell distribution is localized in space, summing these sinks create a local gradient that in turn may trigger a chemotactic response (22). Self-organized gradients naturally arise in bacterial populations consuming nutrients or oxygen, producing the traveling chemotactic bands first described by Adler (23), as well as in eukaryotic systems including starved *Dd* (24), melanoma cells (25), and macrophages (26) through the degradation or uptake of various molecules. In all of these cases, cell migration and modification of the chemical environment form a feedback loop: cells generate or reshape the gradient that subsequently directs their motion. Such feedbacks are naturally described using Patlak-Keller-Segel (PKS) -type continuum models, in which the collective dynamics are governed by effective transport coefficients that incorporate the microscopic behavior of individual cells (27–29). Numerous variants of PKS models have been introduced to model pattern formation, travelling bands and other related collective behaviors (30– 32).

Among self-generated gradients, oxygen (O_2_) occupies a unique position because it is simultaneously a nutrient, a signaling molecule, and a universal determinant of cellular metabolism (33). Although O_2_ sensing and aerotaxis (*i*.*e*.., migration toward O_2_) have long been recognized in bacteria (34, 35), only recently has it become clear that self-generated oxygen gradients can also organize collective migration in eukaryotic cells. A major advance came from the discovery that epithelial cells and cells from sixteen human breast tumour biopsies confined beneath a glass coverslip spontaneously generate oxygen gradients through their own respiration, thereby driving collective aerotactic migration (36, 37). Using the same assay, we subsequently demonstrated that colonies of the social amoeba *Dd* develop a stable propagating ring that persists for days while maintaining an almost constant cell density in the colony interior. Aerotaxis was confirmed through a single-cell microfluidic experiment. The overall spreading behavior was quantitatively explained using a modified PKS-type “Go-or-Grow” model that couples oxygen consumption, aerotaxis and proliferation with a threshold at approximately 1% pO_2_ that triggers aerotaxis below or cell proliferation above it (38). This model not only reproduced wild-type colony dynamics but also provided a quantitative framework for interpreting a large collection of metabolic *Dd* mutants simply through changes in the value of the transport coefficients (cell diffusion, aerotaxis) and reaction coefficients (O_2_ consumption, proliferation), without introducing additional biological mechanisms (39).

These previous studies show that PKS-type descriptions can successfully connect individual responses to environmental cues with collective migration at the population scale. This raises a broader question: how should such macroscopic descriptions be modified when the effective transport properties themselves emerge from interactions within the population? In that case, motility and directional response may no longer be determined solely by the extracellular environment, but may also reflect the local organization and interactions of the cells. Addressing this question requires a system in which transport coefficients themselves become strongly density-dependent. *Acanthamoeba castellanii* (*Ac*) provides such a model. Unlike *Dd, Ac* cells exhibit long persistence lengths that decrease markedly with increasing cell density owing to collision-induced reorientations. Previous work by our group has shown that these microscopic interactions lead to continuous variations in cellular persistence as a function of density, from which the diffusion coefficient can be quantitatively determined using kinetic theory (40). However, we have not investigated any potential modification of the advective transport coefficient (aerotaxis), though it is reasonable to assume that it, too, will be altered. In stark contrast to *Dd, Ac* is a strictly solitary amoeba (41, 42), it does not form multicellular aggregates, lacks coordinated chemical signaling (43), and exhibits negligible cell-cell adhesion, maintaining its independent behavior even in highly crowded environments. *Ac* therefore provides an ideal system to investigate up to very large densities how the collision-controlled emergence of transport coefficients shapes collective aerotactic migration and to test the predictive power of the PKS framework beyond the constant-coefficient regime.

This work addresses two fundamental questions. Can robust collective migration be quantitatively explained when macroscopic transport coefficients emerge from microscopic collision dynamics? And if so, can these collision-dependent transport coefficients predict macroscopic colony dispersal without phenomenological fitting? To answer these questions, we combine the confinement assay with quantitative oxygen mapping using phosphorescent sensing films, oxygen-controlled microfluidics, and single-cell tracking. These independent measurements are integrated into a PKS-type drift-diffusion model whose transport coefficients are entirely determined from experiments and microscopic kinetic theory, without adjustable parameters. Beyond explaining the aerotactic spreading of *Ac*, this framework establishes a quantitative strategy for linking single-cell dynamics to collective migration and for comparing oxygen sensing and aerotactic responses across different motile cell types.

## Results

### Colony spreading provides a simple and quantitative framework for probing the microscopic mechanisms underlying collective cell migration

The expansion of initially confined bacterial or eukaryotic cell colonies has long been used as a macroscopic readout of individual motility, allowing transport properties such as effective diffusion or chemotactic bias to be inferred from population-scale dynamics (7, 38, 44, 45). Here, we investigate the spreading of the axenic free-living strain 30010 of *Acanthamoeba castellanii* using a stencil-based seeding method that enables independent control of the initial colony size and cell density. We begin with the simplest configuration, in which circular colonies spread freely without a coverslip and therefore in the absence of confinement-induced oxygen gradients or other directional cues (Fig. 1A). Individual trajectories recorded over 1.5 h (Fig. 1B) reveal two distinct motility regimes within the same colony. Cells located at the expanding edge follow long, smooth and highly persistent trajectories, whereas cells remaining in the colony interior undergo frequent collision-induced reorientations and consequently display less persistent motion. Despite the net expansion of the colony, cell trajectories exhibit no preferential radial orientation, indicating that spreading in the absence of confinement is driven solely by random motility rather than by a directional migratory bias. These observations are fully consistent with our recent quantitative analysis of random *Ac* motility in homogeneous environments (40), which showed that increasing cell density progressively reduces trajectory persistence and the effective diffusion coefficient through collision-induced reorientations, while leaving the instantaneous migration speed essentially unchanged. The freely spreading colonies shown here therefore provide the reference state from which the effects of self-generated oxygen gradients can be distinguished in the following experiments

**Fig. 1.**
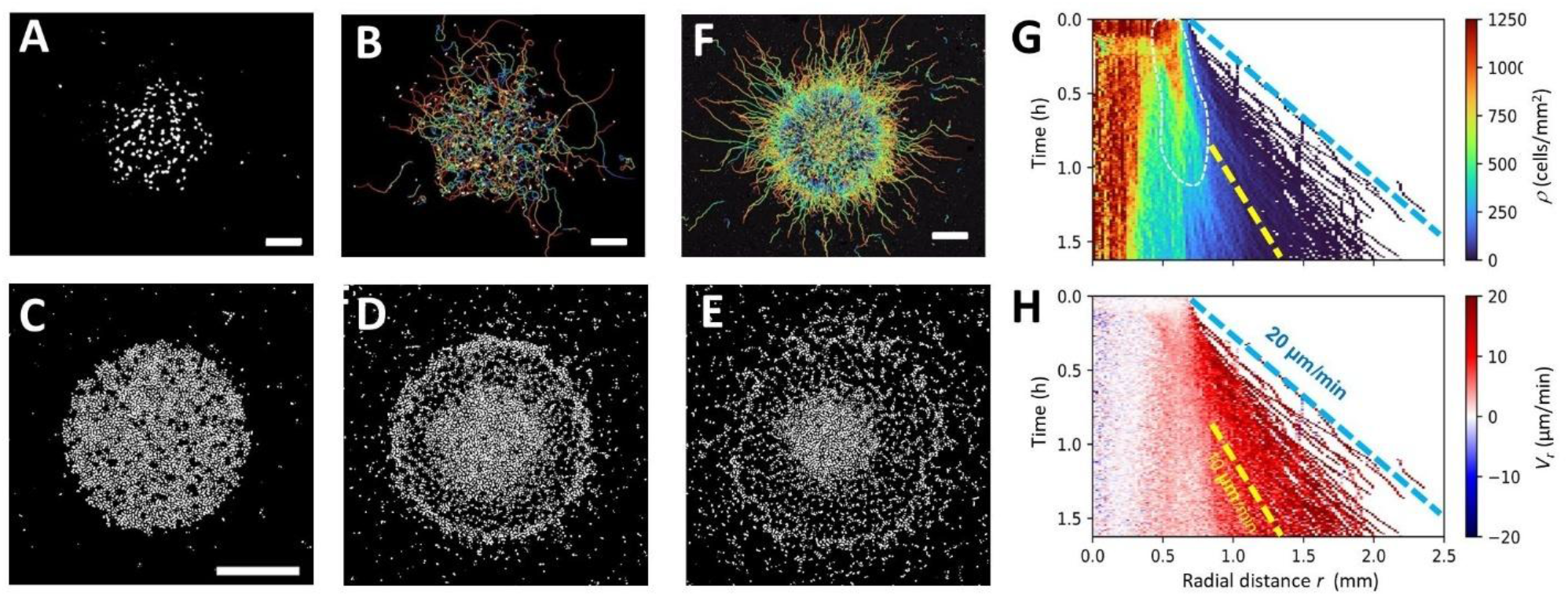
Density-dependent spreading dynamics of *Ac* in unconfined and confined spots. (A) Initial snapshot of an unconfined spot seeded at *ρ* = 90 cells/mm^2^. (B) Cell trajectories over 90 min for the same unconfined spot, showing persistent motion at the periphery and colliding trajectories in the center. (C–E) Snapshots at *t* = 0 (C), *t* = 40 min (D), and *t* = 90 min (E) of a covered spot seeded at high density (*ρ* = 1200 cells/mm^2^), highlighting the four spreading zones: crowded core, intermediate region, diffuse ring, and ballistic cells. (F) Corresponding 80-min trajectories for the high density confined spot. (G-H) Density (G) and radial velocity *V*_*r*_ (H) kymographs for the same confined spot. The region outlined by a white dotted line in (G) corresponds to the dissipating ring. Yellow and blue dotted lines indicate ballistic cell trajectories with radial velocity of 10 µm/min and 20 µm/min respectively. Scale bars, 500 µm.

### Confinement induces ballistic leader cells and a dissipating density band

When a circular *Ac* colony (*ρ* = 1200 cells/mm^2^) was confined beneath a glass coverslip using the assay previously developed for *Dd* **(38)**, its spreading dynamics differed strikingly from those observed in uncovered colonies. Four distinct concentric regions rapidly emerged (Fig. 1C,D and Movie S1). From the center outward, these consisted of (i) a densely packed core displaying highly constrained motion; (ii) a lower-density intermediate region; (iii) a slowly propagating high-density ring; and (iv) a population of rapidly escaping cells at the colony periphery (Fig. 1F). The long, nearly straight trajectories of these escaping cells are reminiscent of ballistic motion. We therefore refer to them as *ballistic cells*, emphasizing their highly persistent migration and their major contribution to colony expansion. Remarkably, the few cells initially located outside the seeded colony first migrated randomly before abruptly switching to persistent outward motion as soon as they encountered the advancing ballistic front (Movie S1), suggesting that they had become exposed to the same self-generated guidance cue.

Cell-density kymographs (two-dimensional plot of the density as a function of the radial coordinate *r* and time *t*, averaged all over angles) revealed the fine details of the slow temporal evolution of these regions (Fig. 1G). The core and intermediate zones progressively decompressed. The ring (demarcated by a white dotted line in Fig. 1G) initially propagated outward at a velocity below 5 µm/min, as determined by the kymograph slope, and eventually dissipated within 1 h. The comet-like trajectories of individual ballistic cells were readily detected in both density and radial velocity kymographs (Fig. 1G-H). In the outer region ahead of the ring, ballistic cells escaping early (t ≃ 0–0.5 h) exhibited higher radial velocities (blue dotted line in Fig. 1I–J, *V*_*r*_∼20 µm/min) than those initiating outward motion at later times (t≃ 1 h, yellow dotted line, *V*_*r*_ ∼10 µm/min).

This complex spreading pattern was independent of both colony geometry and the size or shape of the coverslip, provided the coverslip exceeded the colony dimensions. Spreading from a covered, elongated rectangular stripe of cells provided a simpler, quasi-one-dimensional (1D) Cartesian geometry to analyze spatiotemporal dynamics and associated signals (SI Appendix, Figs. S1, S2 and Fig. 2A–C). Regardless of the coverslip surface (bare glass: SI Appendix, Fig. S1A–C and Movie S2; or glass spin-coated with a PDMS oxygen-sensing film: Fig. 2A–C), high-density rectangular stripes (*ρ*=1200 cells/mm^2^) consistently exhibited the same regions as circular spots during spreading. From the periphery inward, these regions comprised: ballistic cells, a slowly propagating outward band of accumulated cells (analogous to the ring observed in circular geometries), an intermediate zone, and eventually a crowded core. These spatial domains are clearly visible in longitudinal density profiles along the *x-*direction, perpendicular to the long axis of the stripe (SI Appendix, Fig. S1D), as well as in *x-t* density kymographs (SI Appendix, Fig. S1F and Fig. 2E).

**Fig. 2.**
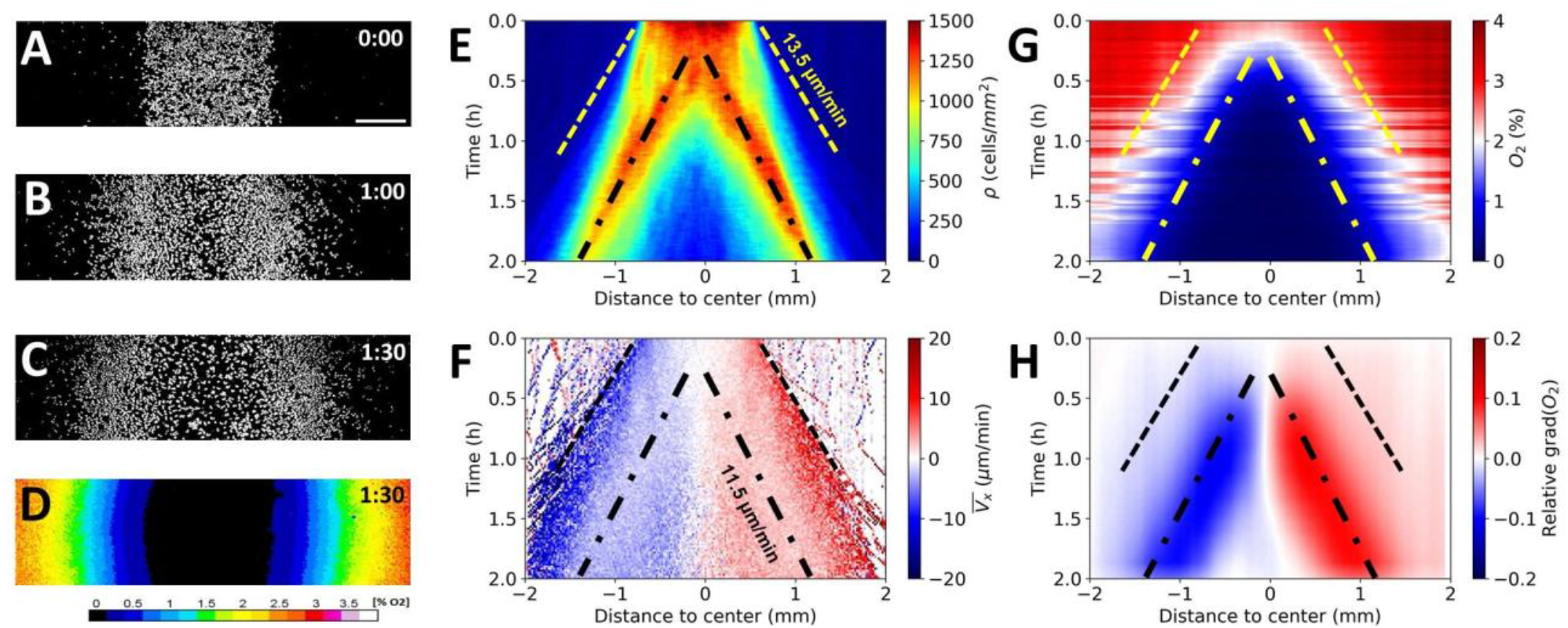
Coupled cell-density and oxygen dynamics reveal aerotaxis-driven spreading in confined *Ac* stripes. (A–C) Time-lapse snapshots at t = 0 (A), 60 min (B), and 90 min (C) of a confined rectangular stripe of *Ac*, showing the outward propagation of an accumulated cell band and fast, outward-directed ballistic cells. The stripe is confined beneath a glass coverslip coated with a ∼50-μm-thick O2-sensing film (PDMS embedding PtTFPP, a porphyrin phosphorescent dye). Scale bar: 500 µm. (D) Reconstructed O_2_ map beneath the coverslip at t = 90 min, obtained from porphyrin phosphorescence and calibrated using the Stern–Volmer relation (color scale indicates O2 concentration *C*). (E-H) Kymographs (*x* position vs. time representation) of cell-density, x-component velocity *V*_*x*_ calculated over a frame interval Δt = 1 min, oxygen *C* and relative oxygen gradient *l* ∇*C*/*C*. All kymographs are averaged along the vertical *y* direction. Outer dashed lines in (E-H) mark the trajectories of the leading ballistic cells, while the inner dash-dotted lines trace the propagating cell band.

Density kymographs obtained from rectangular stripes in Cartesian geometry were qualitatively similar to those in radial geometry (Fig. 1G), with only minor differences in the density band of accumulated cells. Although this band slowly dissipated over longer timescales, as indicated by its decreasing width and contrast (SI Appendix, Fig. S1F and Fig. 2E), it exhibited higher stability than the circular ring, remaining visible beyond 2 h (bare glass) and 1.5 h (PDMS sensing film). This difference likely stems from geometric constraints: in the absence of cell division, the local density of an expanding circular ring must decrease due to radial dilution, whereas a linear band in Cartesian geometry can maintain a constant density as it propagates. Nevertheless, regardless of the geometry, the slope of the band decreased over time, reflecting progressive deceleration. Notably, under the oxygen-sensing film (Fig. 2E), the band emerged later and was wider than under the bare coverslip (SI Appendix, Fig. S1F), likely owing to subtle differences in confinement and the associated oxygen boundary conditions.

Ballistic cells emerged consistently across all experimental configurations. For instance, in striped colonies confined by a bare coverslip, trajectory analysis (SI Appendix, Fig. S1E and Movie S2) and velocity kymographs (*V*_*x*_, *V*_*y*_) revealed that ballistic trajectories were almost perfectly aligned along the *x-*direction. These dynamics exhibited strongly asymmetric expansion with *V*_*x*_≈10–20 µm/min and *V*_*y*_≈0 (SI Appendix, Fig. S1G, H). The longitudinal velocity *V*_*x*_ of individual ballistic cells remained nearly constant throughout their outward escape, as evidenced by the uniform slopes of their trajectories in *x-t* kymographs (SI Appendix, Fig. S1F–H). This constant speed persisted for up to 14 h in large-field-of-view stereomicroscopy experiments, which enabled cell tracking until escape from the confined region beneath the coverslip (SI Appendix, Fig. S2 and Movie S3). Here again, cells initiating ballistic motion at later times displayed similarly constant trajectories but with shallower slopes—reflecting reduced speeds—in both circular (blue vs. yellow slopes in Fig. 1G, H) and linear geometries (Fig. 2E, G and SI Appendix, Figs. S1F, S2D).

### Distinct spreading dynamics in *Ac* and *Dd* under confinement

The spreading pattern of covered *Ac* colonies differs markedly from that observed in *Dd*. An example of propagating *Dd* rings under confinement is shown in (SI Appendix, Fig. S3). In *Dd*, the ring velocity eventually reaches a steady value that remains stable for days (38), and no ballistic cells are observed ahead of the front. Pronounced differences between the two species become particularly evident in the low-density confined regime. In *Dd*, although the time required for ring formation can be long at low initial density (38), rings invariably emerge once cells proliferate sufficiently within the confined region. In contrast, *Ac* cells migrate three-to fourfold faster than *Dd* cells but proliferate much more slowly (doubling time of ∼24 h, compared to ∼8 h for Dd). Consequently, low-density *Dd* colonies first undergo slow local densification prior to ring formation whereas low density *Ac* expand outward but do not form rings. Overall, we identify in *Ac* a critical density threshold of approximately (*ρ*_*c*_≈1100 cells/mm^2^) above which a band of accumulated (corresponding to a ring in circular geometry) emerges. Unlike in *Dd*, however, *Ac* bands are not steady propagating structures: they progressively decelerate and dissipate over time, while ballistic cells are continuously expelled at high, constant velocities.

### The spreading pattern of *Ac* is driven by collective long-range avoidance

We previously showed that *Dd* rings arise from oxygen consumption beneath the colony and propagate along iso-oxygen contours, with a constant wave-speed. The distinct dynamics of *Ac* bands and the presence of ballistic cells raise the question of whether mechanisms beyond oxygen gradients alone are involved. To address this, we designed a stencil-based assay in which a rectangular cell stripe was positioned adjacent to a circular spot, thereby breaking radial symmetry (SI Appendix, Fig. S4 and Movie S4). Cells were seeded at low density (∼100 cells/mm^2^), and we compared unconfined (SI Appendix, Fig. S4A–B) confined conditions (SI Appendix, Fig. S4C–D). Initial migration patterns differed markedly between the two configurations: under confinement, cells from the stripe expanded predominantly along the *x*-direction, whereas cells from the spot expanded radially outward; unconfined cells, by contrast, displayed largely random trajectories. As the two expanding populations approached each other, the trajectories of confined cells were clearly altered by long-range avoidance, remaining highly persistent while bending markedly away from the opposing colony (SI Appendix, Fig. S4D and Movie S4). In the unconfined configuration, trajectories from opposing colonies overlapped and mixed freely, showing no evidence of long-range avoidance beyond local collision-induced reorientations (SI Appendix, Fig. S4B). Thus, confinement induces a guided collective avoidance phenomenon acting over distances far exceeding direct cell–cell contact. Because *Ac* cells are solitary and not known to communicate via secreted signaling molecules (40, 43), aerotaxis emerges as the most plausible mechanism underlying this long-range repulsion.

### Experimental oxygen mapping confirms that confined Ac cells spread by following the self-generated O_2_ gradient

To directly test the relationship between local oxygen fields and observed spreading patterns, we measured oxygen profiles beneath high-density cell stripes (*ρ* = 1500 cells·mm^− 2^) confined beneath coverslips spin-coated with an O2-sensing film. Under these conditions, the stripes developed a propagating band of accumulated cells and fast, outward-escaping ballistic cells (Fig. 2A–C), yielding a characteristic “tooth-shaped” density kymograph (Fig. 2E). The propagation speed of the cell band was approximately 11.5 µm/min, as estimated by the slope of the dashed-dotted lines obtained from the density kymographs. In contrast, individual cell tracking within this band yielded lower speeds while the speed obtained from the tracks of individual cells in this band region was lower (∼5 µm/min in *Vx* velocity kymographs; Fig. 2F), suggesting local jamming behavior. The first ballistic cells escaping the initial stripe exhibited outward velocity of ∼13.5µm/min (dashed lines in Fig. 2E-H). Oxygen mapping revealed a strongly hypoxic central zone (*C*< 0.4% pO_2_) flanked by a steep, self-generated gradient co-aligned with the direction of cell expansion (Fig. 2D and Movie S5). Spatiotemporal analysis *x-t* O2 kymograph (Fig. 2G) demonstrated that ballistic cells initiated directed motion near the 2% pO2 isoline (dashed line) whereas the cell band propagated within regions where oxygen levels fluctuated around 1% pO_2_ (dashed-dotted lines), corresponding to the region of maximum relative logarithmic oxygen gradient *l∇C/C* at about 0.1 (Fig. 2H) where *l=30µm* corresponds to the characteristic cell diameter of *Ac* cells. The choice of computing this quantity is motivated by the fact we previously showed that logarithmic sensing proportional to *∇C/C* provides a more accurate description of aerotaxis than direct gradient sensing *∇C* (46). *l∇C/C* corresponds to the relative oxygen difference between front and back of *Ac*. These findings directly support an aerotactic response driven by self-generated hypoxia. Interestingly, isolated background cells outside the main front initially moved randomly without directional bias at local O_2_ levels above 3% (Movie S6). However, as leader cells approached and depleted local oxygen, these background cells aligned their migration with the outward stream, accelerating to comparable speeds. Altogether, these results demonstrate that the complex spreading pattern of confined *Ac*, comprising a dissipating cell band and fast, outward-directed leader cells, arises from aerotaxis, with directional migration being activated below 2% O_2_ and robustly maintained across space.

### Aerotactic in *Ac* is quantitatively controlled by logarithmic oxygen sensing with a sharp high-oxygen cutoff

To quantify the precise dependence of the aerotactic displacements on local oxygen gradients, we used a two-layer microfluidic device generating stable and tunable O_2_ profiles, as previously described (38, 46). In this setup (Fig. 3A), two gas channels delivering controlled gas mixtures are positioned above three liquid media channels in which *Ac* cells migrate (Fig. 3B). This superimposed geometry establishes a stationary O_2_ profile in the lower layer, allowing precise control over the local O_2_ concentration experienced by the cells. The O_2_ profile was measured previously (38) and showed good agreement with COMSOL Multiphysics simulations (Fig. 3C, red line, see Methods). The concentration *C* spans from 0.4% to 21% pO_2_ along the *x*-direction when pure N_2_ and ambient air are supplied to the left and right gas channels, respectively. We also computed the dimensionless relative O_2_ gradient *l∇C/C(x)* (Fig. 3C, black line). At the most hypoxic location at *x*=−1.6 mm from the center, the slope of the O_2_ concentration gradient inverted (dotted line in Fig. 3C). Alternative gradient profiles were also evaluated, each showing a gradient inversion position around −1.6 mm (Fig. 3D, F). In the case of the 0-2% gradient for example, soon after gradient establishment, cells migrated away from the inversion position (Fig. 3B, green arrows and green dotted line) creating a pronounced depletion region there at 81 min.

**Fig. 3.**
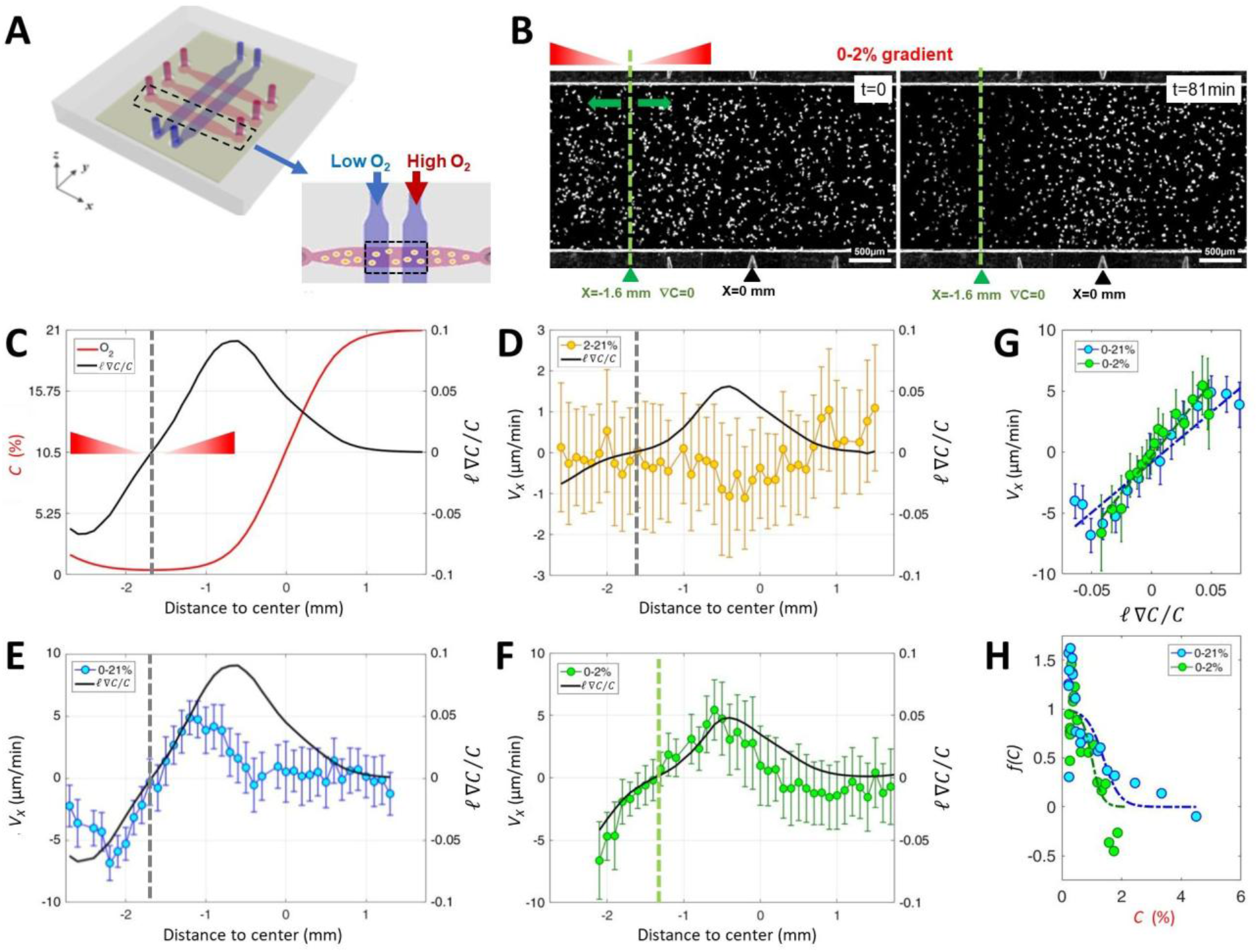
Quantifying the logarithmic aerotactic response of *A. castellanii* using a two-layer microfluidic oxygen gradient generator. **(A)** Schematic of the two-layer microfluidic device used to measure the aerotactic index of *A. castellanii*. Two gas channels (blue) delivering given O_2_ concentrations are positioned above three media channels containing migrating cells (pink), generating a stable oxygen gradient in the lower layer. **(B)** Snapshots of cell distribution in the microfluidic device at initial time t=0 (left) and at t=81 min (right panel) after gradient establishment; cells migrate either to the right or to the left from the green dotted line where the O_2_ concentration is minimal and the oxygen gradient *∇C* is zero. **(C)** Computed oxygen concentration profile *C*(*x*) (red line) and relative dimensionless gradient *l* ∇*C*/*C*(*x*) (black line), where *l*=30µm is the cell diameter, as a function of position *x* along the media channel obtained using COMSOL Multiphysics. **(D–F)** Mean aerotactic velocity *V*_*x*_ (bullets) and *l* ∇*C*/*C* (black line) as a function of *x* for three imposed oxygen profiles: 0–2%, 0-21% and 0–2% (yellow, blue and green bullet points respectively). The gray or green dotted lines in **(C–F)** correspond to the positions at which ∇*C* vanishes. **(G)** Aerotactic velocity *V*_*x*_ as a function of *l* ∇*C*/*C* for low oxygen levels (C<1%). The linear fit yields the aerotactic coefficient *χ*. **(H)** Ratio *V*_*x*_ / *l* ∇*C*/*C* plotted as a function of oxygen concentration *C* revealing the sigmoidal cutoff function *f(C)* that suppresses aerotactic sensitivity above a critical concentration *C*^∗^. In **(G–H)** blue and green bullet points correspond to the 0-21% and 0–2% gradient conditions respectively.

By tracking cell trajectories, we computed at each position *x* along the media channel the cellular flux *ρV*_*x*_, defined as the product of the local cell density *ρ* and the mean drift velocity component *V*_*x*_, along the oxygen gradient. When spatial density gradients exist, a fraction of this flux originates from passive diffusion described by Fick’s law (-*D∇ρ*, where *D* is the cell diffusion coefficient). However, this diffusive contribution is more than one order of magnitude smaller than the measured flux *ρV*_*x*_ in presence of an O2 gradient. Moreover, in the absence of any imposed O2 gradient, *ρV*_*x*_ ≈0, confirming that the measured net flux is dominated by aerotaxis rather than by diffusion. We hence systematically compared *ρV*_*x*_ to the relative oxygen gradient *l∇C/C(x)* for different imposed oxygen profiles (Fig. 3D–F).

Under a 0–21% imposed gradient, *V*_*x*_ closely followed *l∇C/C* over positions extending from x∼-2.2 mm to x∼-1.1 mm. This region corresponds to O_2_ ≤0.75% pO_2_ and encompasses both the minimum and maximum values of *V*_*x*_. Beyond this range, the velocity rapidly collapsed to zero despite the persistence of a finite gradient (Fig. 3E). Under a 2–21% imposed gradient (Fig. 3D), *Ac* cells exhibited no measurable aerotaxis: *V*_*x*_ fluctuated around zero despite the presence of a non-zero logarithmic gradient. In contrast, under a 0–2% gradient (effective profile between 0.35% and 2.35% pO_2_), aerotactic drift remained robust and strictly proportional to *l∇C/C* throughout the channel (Fig. 3F). Together, these observations demonstrate that *Ac* cells become effectively blind to oxygen gradients above a sharp threshold of approximately 2 % pO_2_, whereas below this threshold, their drift velocity is directly proportional to the logarithmic gradient *l∇C/C*.

To quantitatively validate logarithmic oxygen sensing, we plotted *V*_*x*_ as a function of *l∇C/C* for low oxygen concentrations (C≤0.75%, Fig. 3G). Linear fits yielded aerotactic bias speeds *X*=82±15 and 121±18 µm/min for the 0–21% and 0–2% gradient conditions, respectively (95% confidence intervals of the fits). Having established this proportionality regime, we defined an oxygen-sensitivity cutoff function *f(C)* as:

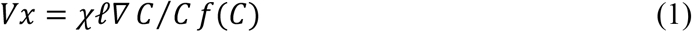

which characterizes the dependence of the aerotactic response on oxygen concentration. The function was fitted using a sigmoid expression:

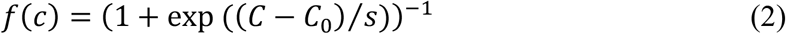

where *C*_*0*_ is the oxygen concentration at which the aerotactic response is reduced by half and *s* is a steepness parameter. Experimentally we measured *C*_*0*_=1.4±0.5% and 1.0±0.4%, *s*=0.3±0.4% and 0.2±0.3% for 0–21% and 0–2% conditions, respectively (Fig. 3H).

### The aerotactic bias and diffusion coefficient decrease with cell density, whereas the oxygen sensitivity cutoff remains unchanged

We next examined how the key aerotactic parameters *X, C*_*0*_ and *s* vary with cell density *ρ* (Fig. 4A–C). The two gradient conditions discussed above corresponded to relatively similar densities (*ρ*=142 and 92 cells/mm^2^, respectively), but a broader systematic analysis revealed a pronounced decrease of the aerotactic bias speed with increasing density (Fig. 4A). For comparison, we also considered in Fig. 4D the density dependence of the diffusion coefficient *D*(*ρ*) previously obtained from analyses of random *Ac* trajectories in isotropic normoxic environments (40). In our previous work, the effect of cell-cell collisions was described at a mean-field level through a density-dependent reorientation rate, or an inverse persistence time, *λ*(*ρ*) = *λ*_0_ + *αρ* where *λ*_0_ is the intrinsic reorientation rate in the absence of collision, leading to *D*(*ρ*) ∝ (*λ*_0_ + *αρ*)^−1^. Motivated by the similar decrease observed for the aerotactic bias, we introduced the phenomenological form

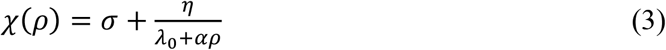

**Fig. 4.**
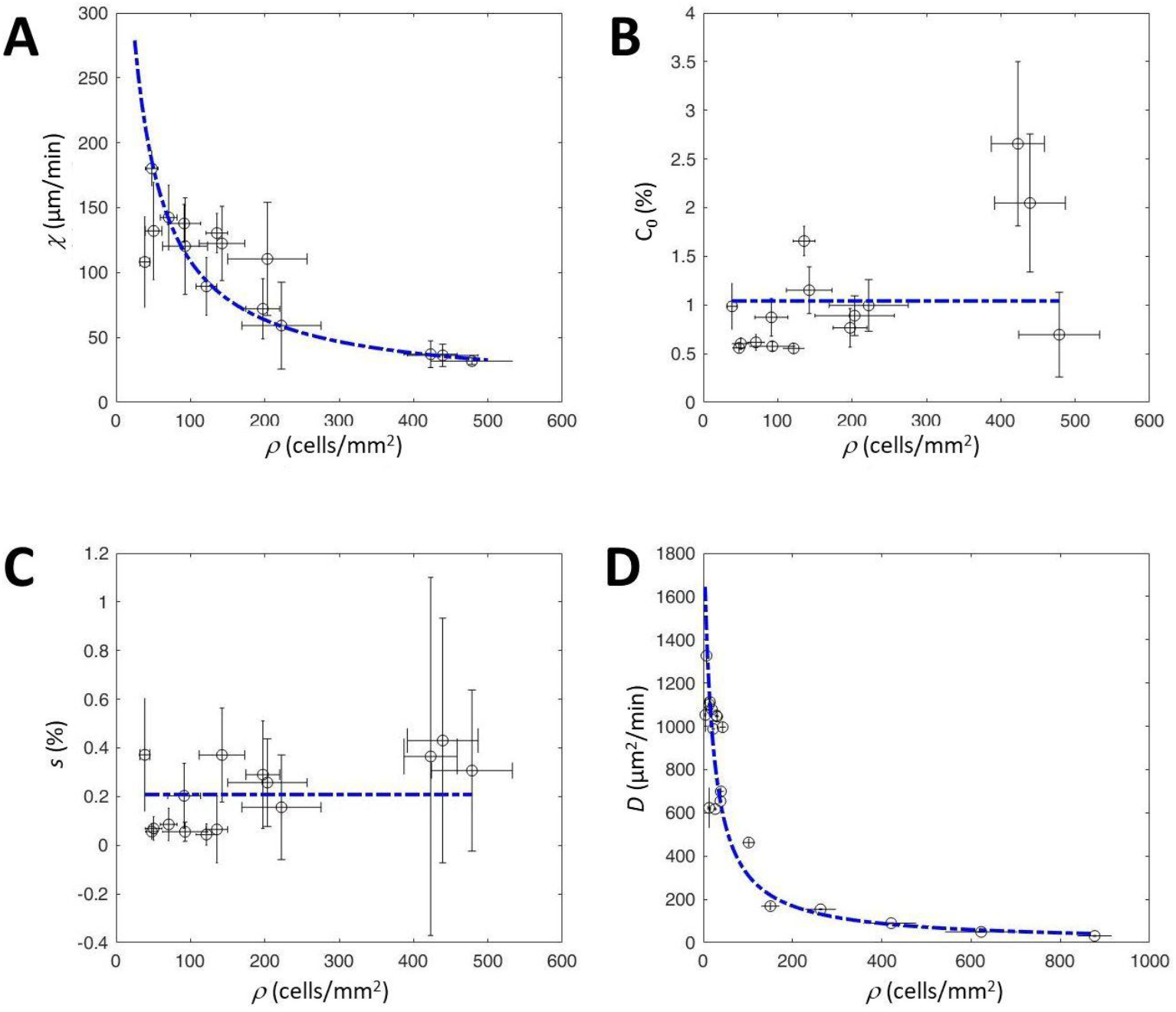
*Ac* aerotactic and single cell diffusion parameters at increasing cell densities. **(A)** aerotactic bias speed *X*, **(B)** cutoff *C*_*0*_ in O_2_ concentration at which the aerotactic response relative to gradient is diminished by a factor of two. **(C)** Steepness parameter s defined as the width of the sigmoidal cutoff function. **(D)** diffusion coefficient *D* obtained from random cell trajectories in isotropic and normoxic environments. The vertical error bars represent the 95% confidence intervals obtained from fits discussed in the text. The horizontal errors bars represent std of *ρ* obtained at different location in the device. The blue dotted lines in (A), (D) represent fits with the form *χ*(*ρ*) = *η*/(*ρ* + *β*) and *D* = (*αρ* + *δ*)^−1^. The blue dotted lines in (B), (C) represent mean values over the full range of *ρ*. All parameter values *η, β, α, δ*, 〈*C*_0_〉 and 〈*s*〉 are indicated in the main text.

Below, we propose a microscopic rationale for this phenomenological form.

The diffusion and aerotactic measurements were then fitted simultaneously, using the same density-dependent reorientation rate *λ*(*ρ*) = *λ*_0_ + *αρ* in both coefficients. To constrain the parameters, we fixed the value of *α* to the experimental collision rate *α*_*C*_ = (10.5 ± 1.2) × 10^−4^ min^-1^·mm^2^·cell^-1^ measured in Ghazi et al. (40). For this joint fit of 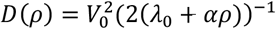 and *χ*(*ρ*) = *σ* + *η*(*λ*_0_ + *αρ*)^−1^, we find *V*_0_ = 8.2 ± 1.7 µm·min^-1^, *λ*_0_ = 0.019 ± 0.008 min^-1^, *σ* = 10.0 ± 5.2 µm·min^-1^ and *η* = 12.3 ± 2.9 µm·mm^-2^. In contrast, the cutoff parameters *C*_0_ and *s* showed no significant dependence on cell density, with average values *C*_0_=1.04±0.6 % and *s* =0.21±0.14% (Fig. 4B-C).

Together, these results demonstrate that the aerotactic response of *Ac* cells is governed by logarithmic oxygen sensing with a sharp cutoff above *C*_*0*_ ≈1%, largely independent of cell density. Below this threshold, however, the aerotactic bias strength *χ* decreases strongly with increasing density, similarly to the collision-controlled reduction of the effective diffusion coefficient *D*. We now provide a possible microscopic rationale for the phenomenological density dependence of *χ*, based on collision-induced changes in reorientation frequency.

### Collision-controlled directional persistence provides a microscopic rationale for density-dependent aerotaxis

Before incorporating these experimentally determined transport coefficients *D*(*ρ*) and *χ*(*ρ*) into a PKS-type “go or grow” model, we briefly discuss a possible microscopic rationale for the density dependence of *χ*(*ρ*). In our previous work on random *Ac* migration in isotropic environments (35), we described the effect of cell-cell collisions using a mean-free-path approach. The reorientation rate, which is the inverse of the characteristic persistence time, was written as *λ*(*ρ*) = *λ*_0_ + *αρ*, where *λ*_0_ is the inverse of the characteristic persistence time in the absence of collisions and *αρ* accounts for the additional reorientations induced by cell-cell collisions. As cell density increases, the persistence time *λ*(*ρ*)^−1^ decreases, leading to the observed decrease of the diffusion coefficient *D*(*ρ*). Here, we ask whether the same collision-dependent reorientation rate can also provide a microscopic rationale for the density dependence of aerotactic motion. In a minimal one-dimensional framework, cells move with velocity *V*_0_ either toward (+ ) or away (− ) from higher oxygen concentrations. Aerotaxis can then arise through two complementary microscopic mechanisms, both proportional to the filtered logarithmic oxygen signal *ϕ*[*C*] = *f*(*C*)*l*∇*C*/*C*.

First, cells moving in the favorable direction may reorient less frequently than cells moving toward lower oxygen concentrations, introducing a temporal bias in the reorientation dynamics. We describe this by setting *λ*_+_ = *λ*(*ρ*) − *κϕ*[*C*] and *λ*_−_ = *λ*(*ρ*) + *κϕ*[*C*], where *κ* > 0 measures the strength of the temporal bias. Such modulation of turning frequency is conceptually analogous to bacterial chemotaxis, where runs oriented toward favorable conditions are prolonged whereas unfavorable runs are shortened (16, 30). Because this mechanism directly acts through the reorientation frequency, it is naturally affected by collision-induced reorientations and therefore decreases with increasing density.

Second, after a reorientation event, the direction selected may itself be biased toward higher oxygen concentrations, introducing a spatial bias. We describe this by setting the probability of selecting the positive (resp. negative) direction *p*_+_ = 0.5 + *γϕ*[*C*] (resp. *p*_−_ = 0.5 − *γϕ*[*C*]), where *γ* > 0 measures the strength of the spatial bias. This mechanism is reminiscent of spatial gradient sensing in eukaryotic chemotaxis, where spatial differences in the sensed chemoattractant bias migration toward higher concentrations (17). In contrast to the temporal bias, this contribution acts through the direction selected after a reorientation rather than through the persistence time. Consequently, if collisions alter the reorientation frequency without affecting this directional choice, its contribution to the aerotactic drift remains approximately independent of cell density.

The stationary probabilities of motion in each direction can be computed analytically using this two-state persistent random-walk description (see SI appendix for detailed calculations). Neglecting higher-order corrections in the aerotactic signal, whose magnitude is assessed in the SI appendix, the resulting aerotactic drift velocity takes the form

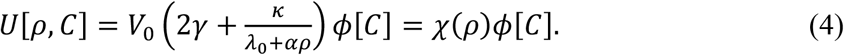

Hence by comparison with the empirical Eq. (3), *σ* = 2*V*_0_*γ* and *η* = *V*_0_*κ*.

The density-independent term *σ* of *χ*(*ρ*) corresponds to the spatial-bias contribution, whereas the term *η*/(*λ*_0_ + *αρ*) originates from the temporal bias and decreases as the collision-induced reorientation rate increases. A closely related density dependence of the chemotactic coefficient has been reported for dense populations of *E. coli* by Phan *et al*. (32), where a similar phenomenological form was introduced. Related work by Colin *et al*. (47) showed that hydrodynamically driven collective reorientations at high cell density reduce directional persistence and chemotactic efficiency. This simple construction therefore provides a possible microscopic rationale for the phenomenological form of *χ*(*ρ*) used above. In particular, the same density-dependent reorientation rate, *λ*(*ρ*) = *λ*_0_ + *αρ*, that controls the diffusion coefficient *D*(*ρ*) also enters the density-dependent contribution to the aerotactic coefficient *χ*(*ρ*), providing a possible common mechanism through which cell-cell collisions modulate both random and biased cell motion.

### Collision-controlled transport coefficients reproduce Ac spreading dynamics

To determine whether the experimentally measured density dependences of the diffusion coefficient *D(ρ*) and aerotactic coefficient *χ(ρ)* are sufficient to explain the spreading dynamics of *Ac*, we extended the reaction-advection-diffusion model previously developed for confined *Dd* colonies evolving under self-generated oxygen gradients (38). The model couples the evolution of cell density *ρ* and oxygen concentration *C* through two partial differential equations accounting for random cell motion, aerotactic drift, cell proliferation, oxygen diffusion and oxygen consumption:

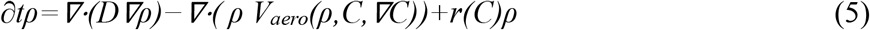

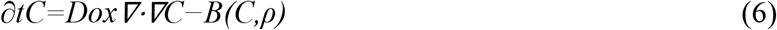

where *V*_*aero*_*(ρ,C,∇C)* is the aerotactic biased velocity, *r* and *B* are the proliferation rate (Fig. S5A) and O_2_ decay rate respectively. The O_2_ decay rate is due to the oxygen uptake by cells located at the bottom of the confinement layer of thickness *h*, hence *B(C)=ρQ(C)/h* where *Q(C)* is the oxygen consumption rate per cell. The model further incorporated a parsimonious “Go-or-Grow” hypothesis: cells proliferate but do not exhibit aerotaxis above a critical oxygen concentration *C*_*Lim*_, whereas below this threshold they become aerotactic and temporarily stop dividing.

In *Dd*, this framework successfully reproduced the long-lived propagating ring (Fig. S3) using constant transport coefficients because cell trajectories are only weakly affected by cell density. We therefore asked whether the same model, still assuming constant transport coefficients and using the aerotactic law *V*_*aero*_ = *χl*∇ *C*/*C f*(*C*), could also account for the markedly different spreading dynamics of *Ac*. Compared with *Dd, Ac* cells migrate considerably faster and consume oxygen approximately three times more rapidly (*Q*_*AC*_=1.3×10^−16^ mol cell^−1^ s^−1^; supplementary Fig. S5B-C), whereas their proliferation rate is approximately threefold lower (*r*_*0*_=ln2/1440 min^−1^ versus ln2/480 min^−1^ for *Dd*; supplementary Fig. S5A). Consequently, proliferation has little influence over the first several hours of spreading and was retained only to simulate longer times.

(whether the high transport coefficients experimentally measured at low density in *Ac* (*D* = *D*0 = 470 µm^2^/min, *χ* = *χ*0 = 175 µm/min), while remaining constant throughout the simulation, could account for striped colony spreading. This condition produced a rapidly expanding cell-density band (peak of density in Fig. 5A,B) following the self-generated 1% pO2 isoline (Fig. 5C), but failed to generate the dense band of cell accumulation propagating behind the front observed experimentally. The peak initially propagated at very high speed but progressively slowed to 22 µm/min after 1 h (Fig. 5C,D). Its shape was initially sharp and progressively lost both height and sharpness, while remaining localized as a density peak (Fig. 5A). Conversely, using constant low transport coefficients measured at high density (*D* = 72 µm^2^/min, *χ* = 37 µm/min) generated a much slower front consisting of a density peak that propagated together with the 1% pO2 isoline at approximately 9 µm/min (SI Appendix, Fig. S6). The peak was initially sharp but rapidly dissipated. Neither set of constant transport coefficients therefore reproduced the characteristic coexistence observed experimentally between a fast ballistic front and a slowly propagating accumulation band. Interestingly, the high-*D*, high-*χ* condition nevertheless produces a sharp density peak at the leading edge, qualitatively resembling the propagating ring observed in *Dd*, although in *Dd* the peak remains much more stable over the experimental timescale because cell proliferation substantially contributes to maintaining the ring structure.

**Fig. 5.**
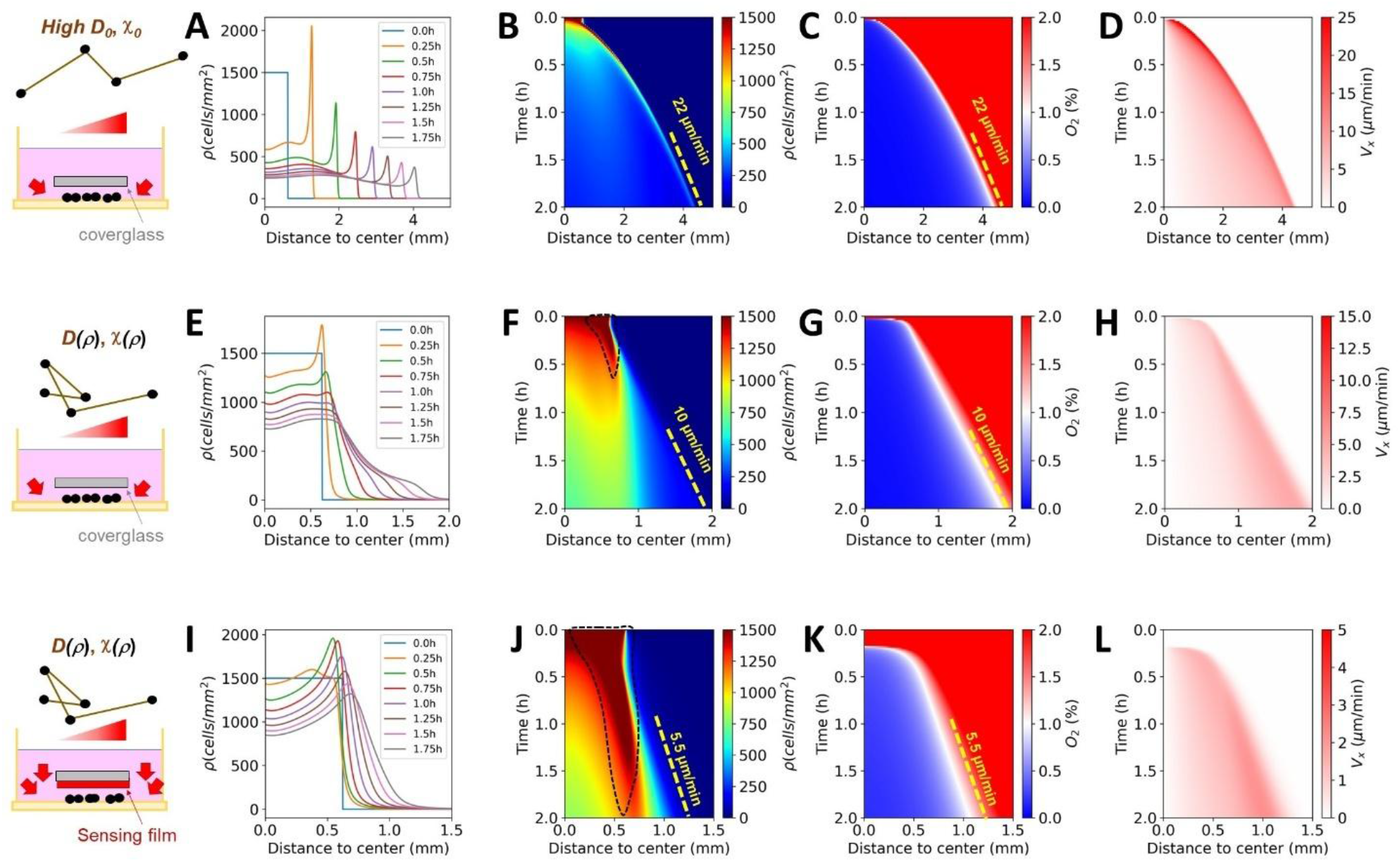
Modelling Ac spreading in cartesian geometry from an infinite confined stripe of cells of width 1.25mm with different transport and reaction coefficients. (A-D) High constant cellular diffusion coefficient (*D*_*0*_=470 µm^2^/min), high constant aerotactic bias speed *X*_*0*_=175 µm/min. (E-H) Density-dependent transport coefficient *D*^−1^(*ρ*) = *ερ* + *δ, χ*(*ρ*) = *σ* + *η*/(*ρ* + *β*) with *ε* = 27 × 10^−6^ min×mm^2^×µm^-2^, *δ* = 5 × 10^−4^ min×µm^-2^, *σ* = 10 µm×min^-1^, *η* = 1.17 × 10^4^ µm×min^-1^×mm^2^ and *β* = 18.5 mm^-2^. (I-L) Same density-dependent transport coefficient but more O_2_ leakages due to the presence of a PDMS based sensing film (this situation is simulated with a lower O_2_ consumption as compared to E-H). (A,E,I) Density profiles as a function of the distance to center in the x direction, (B,F,J) kymographs of oxygen level, (C,G,K) kymographs of cell density, (C,F) and (D,H,L) kymographs of *V*_*X*_ velocity.

Introducing the experimentally measured density dependence of both transport coefficients immediately resolves this discrepancy. The simulations simultaneously generate a frontal zone populated by fast-moving cells, analogous to ballistic cells (Fig. 5E-F, H), that follow the 1% pO2 isoline at a speed of approximately 12 µm/min (Fig. 5G), and behind them, a slowly moving, dissipating band of cell accumulation (see the dashed area in Fig. 5F).

To further challenge the model, we simulated the experiments performed in the presence of the oxygen sensing film (Fig. 5I-L). In this configuration, the PDMS layer acts as an additional oxygen reservoir because oxygen is 6.5-fold more soluble in PDMS than in the PYG90 culture medium (38). Consequently, the effective oxygen depletion term in Eq. (6) becomes *B’(C)=ρQ(C)/h’* where *h*^’^ = *h*_*HL*5_ + *h*_*PDMS*_ × *S*_*PDMS*_/*S*_*HL*5_ with *h*_*PDMS*_ ≈ 25µm and *h*_*HL*5_ ≈ 50µm, thereby reducing the apparent oxygen consumption rate by a factor of about 4.5. Without modifying any cellular transport coefficient, this single physical change decreases the propagation speed of the oxygen depletion front to approximately 5.5 µm/min (Fig. 5K), resulting in a correspondingly slower ballistic front (Fig. 5J). Although lower in amplitude, the predicted reduction closely matches the experimental front velocities reduction measured in the absence or presence of sensing film (20 and 13.5 µm/min, respectively; Fig. S1F and Fig. 2E). Because fewer ballistic cells escape from the initial stripe, cell crowding persists for longer near the colony edge, producing a more pronounced and longer-lived accumulation band (compare the dashed regions in Fig. 5F and Fig. 5J). These predictions again closely reproduce the experimental kymographs.

Together, these simulations demonstrate that the experimentally measured density dependence of the transport coefficients is the essential ingredient required to quantitatively reproduce *Ac* colony spreading. Once incorporated into the PKS framework, these collision-controlled transport coefficients predict the spatial separation between a fast aerotactic front of ballistic leader cells tracking the 1% O_2_ isoline and a slower, dissipative accumulation band remaining behind it. This decoupling, absent in *Dictyostelium* and in simulations with constant transport coefficients where the propagating front and density peak coincide, emerges naturally from the density dependence of the transport coefficients. The same framework also quantitatively predicts how both structures are modified when oxygen transport is altered by the PDMS sensing film.

## Discussion

The present work shows that the collective spreading of confined *Acanthamoeba castellanii* colonies can be quantitatively understood from a remarkably simple physical principle: cell-cell collisions progressively reduce both random motility and aerotactic transport. By combining quantitative single-cell measurements, controlled oxygen microfluidics and reaction-advection-diffusion modelling, we demonstrate that the experimentally measured density dependences of the transport coefficients *D*(*ρ*) and *χ*(*ρ*) are sufficient to predict the coexistence of two spatially distinct structures: a fast aerotactic front composed of ballistic leader cells following the 1% oxygen isoline, and a slower dissipative band of accumulated cells trailing behind. The model further predicts how both structures are modified by the presence of an oxygen sensing film, without introducing any additional adjustable transport parameters.

A major outcome of this work is that the apparently distinct collective behaviours of *Acanthamoeba* and *Dictyostelium* can be understood within the same PKS framework. In both organisms, oxygen consumption generates self-produced oxygen gradients that drive collective aerotaxis. The difference does not arise from distinct sensing strategies or additional biological ingredients, but from the transport coefficients themselves. In *Dictyostelium*, collisions have little influence on the short persistence of cell trajectories, resulting in nearly constant transport coefficients and the propagation of a stable ring in which the migration front and density peak remain superimposed. In contrast, the much larger persistence of *Acanthamoeba* trajectories makes both transport coefficients strongly collision-dependent. This density dependence causes the aerotactic front to detach from the density maximum, producing the characteristic separation between ballistic leader cells and the dissipative accumulation band.

Continuum descriptions of chemotaxis generally treat transport coefficients as phenomenological parameters, sometimes allowing them to vary with cell density to reproduce experiments. A striking recent example is the work of Phan et al., who demonstrated that the classical PKS model breaks down at high bacterial density and restored its predictive power by introducing density-dependent chemotactic transport and nutrient consumption (32). Our work reaches a similar conclusion from the opposite direction. Rather than postulating density-dependent coefficients, we derive them from microscopic collision statistics and independently measure every transport coefficient before confronting the model with experiments. This progression from phenomenological to predictive coarse-graining provides a quantitative link between microscopic cell dynamics and continuum transport equations. Similar coarse-graining strategies underlie kinetic theories of chemotaxis and dispersal in biology (11–13), hydrodynamic theories of active matter (48), and, more recently, continuum descriptions of dense human crowds (49).

Several limitations nevertheless persist within the current framework, which may account for the quantitative discrepancies observed between modeling and experimental results, particularly regarding the width and propagation speed of the accumulation band (Fig; 2E and 5J). We suspect that these deviations stem partly from the inherent differences between the discrete Run-and-Tumble (RT) approach used here and more continuous frameworks like Active Brownian Particles (ABP), even though both classes of persistent random walks become rigorously equivalent at long times in terms of their asymptotic Mean Squared Displacement (MSD) and effective diffusion (50). While our RT approach coupled with a mean free path theory successfully captures the density-dependence of the diffusion coefficient via cell–cell collisions (40), assuming strictly straight trajectories between tumbles remains a coarse approximation for biological cells that inherently experience continuous fluctuations, including translational and rotational diffusion, even when polarized. In parallel, real intercellular collisions frequently obey sophisticated behavioral rules, such as CIL (6) or CEL (7); yet, while density effects do not appear to impact the aerotactic spreading of confined spots in *Dd*, where diffusion is primarily regulated by secreted factors rather than collisions (unpublished), we have not observed any significant post-collision CIL or CEL signatures in *Ac* trajectories either (unpublished). Consequently, further analyses and experiments utilizing novel microfluidic or geometric environments will be necessary to comprehensively resolve these intricate microscopic behaviors.

Ultimately, this study utilizing finely tuned microfluidic devices elucidated the mechanism of oxygen sensing. We first demonstrated that cells detect oxygen logarithmically. A more intriguing outcome concerned the mechanism of oxygen-based guidance. Although *Ac* is a large eukaryotic cell capable of spatial gradient sensing in principle (17), our experimental determination of the *χ(ρ)* indicate that aerotaxis is primarily governed by the temporal component of oxygen sensing (term *η*/(*ρ* + β) in Eq. (3)) whereas the spatial component (term *σ* in Eq. (3)) contributes little to the observed migration (Fig. 4A). In this respect, *Ac* behaves more like motile bacteria, which rely mostly on temporal comparisons because of their small size (16). Surprisingly, this result cannot simply be attributed to exceptionally shallow gradients. The measured relative gradient, *l∇C/C*, reaches values of 0.05 and locally approaches 0.1 (Fig. 3C–F), corresponding to a 5–10% difference in oxygen concentration between the front and rear of a cell. Such differences are well within the range that would appear detectable by a spatial-sensing mechanism (15). However, unlike classical chemoattractants, oxygen is not detected through a known membrane receptor. Instead, it readily diffuses across lipid membranes, and the molecular mechanisms underlying O_2_ sensing remain largely unknown (39). Under these conditions, temporal sensing may represent a more natural strategy than spatial receptor comparison. Our results therefore raise the possibility that temporal sensing is not merely a bacterial adaptation to small cell size but may constitute a more general mechanism of aerotaxis shared by diverse eukaryotic cells, including *Acanthamoeba castellani* and perhaps other oxygen-responsive cell types.

## Materials and Methods

### Cell culture and cell line

The axenic free-living strain *Acanthamoeba castellanii* (*Ac*), ATCC NEF 30010 isolated from soil in the United States; California; Pacific Grove, was used throughout this study. Cells were grown in proteose peptone-yeast-glucose (PYG90) medium (51) as monolayers in 75 cm^2^ tissue culture flasks at 25°C. Prior to each experiment, adherent cells were detached and resuspended at 4 × 10^6^ cells/mL.

### Confinement assays

To generate reproducible initial colonies, we developed a stencil-based confinement assay. For the circular spot stencil, a circular hole was punched at the center of a commercial polydimethylsiloxane (PDMS) sheet with 250 μm thickness (SiMPore, USA) using a biopsy puncher with 1.25 mm diameter. Rectangular stencils (10 × 1.25 mm^2^) were fabricated by punching two adjacent circular holes and cutting the intervening PDMS with a scalpel guided by a ruler. The PDMS stencil thus cut was then carefully positioned into the center of a 35mm diameter well using tweezers. Approximately 100 μL of medium containing the desired concentration of cells was then added to the well, with the cells suspended uniformly within the medium. The plate was incubated for 30 minutes, allowing cells to settle and adhere within the stencil opening. After incubation, the volume of medium was aspirated and the PDMS stencil was gently removed, leaving behind a controlled spot or stripe of adhered cells at the center of the well. 2 mL of fresh medium were added to the well, and a glass coverslip was placed over the spot of cells to confine the colony.

### Oxygen sensing film preparation and calibration

Films were prepared by dissolving 17 mg of 5,10,15,20-Tetrakis-(2,3,4,5,6-pentafluorophenyl)-porphyrin-Pt(II) (PtTFPP, Por-Lab, Scharbeutz, Germany) in 3 mL acetone, mixed with a 4:1 PDMS:curing agent solution and degassed. The mixture was spin-coated onto coverslipes and cured at 60°C for 2h. The final film had a dye concentration of 4 mmol/L and a thickness of 25 μm, measured using a ContourGT-K 3D profilometer (Bruker, USA) after surgical blade sectioning. Films were stored in the dark before their use in fluorescence microscopy. The oxygen concentration *C* measured in our various assays was obtained from the non-linear Stern-Volmer Relationship 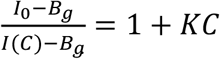 where *I*_0_ is the fluorescence intensity under pure N_2_, *I*(*C*) is the fluorescence intensity for a concentration *C* and *B*_*g*_ is the background. Calibration of the film sensitivity *K* (Stern-Volmer constant) and *B*_*g*_was performed placing the sensing film in a homemade chamber under controlled gas atmospheres. The full image quantification pipeline is detailed in Cochet-Escartin et al (38).

### Microfluidic Devices

The oxygen-controllable double-layer microfluidic device (Fig. 3A) was described previously (38, 52). Briefly, the device is a 30×30 mm square in PDMS (Sylgard 184 Silicone Elastomer Kit, The Dow Chemical Company, USA) with a 4 mm thickness. Inside the device, two gas channels in a 1-mm interval were located at 0.5 mm from the bottom surface, perpendicularly crossing above three parallel medium channels on the bottom. The width and height were 2 mm and 0.15 mm for all channels, respectively. A polycarbonate (PC) film (20 × 25 mm^2^) of low gas permeability was placed on the PDMS layer of the gas channels to prevent oxygen infusion from the atmosphere. The O_2_ concentration in the device was controlled by gas exchange between the channels, supplying gas mixtures at predefined oxygen concentrations to the gas channels. The oxygen concentration in the device was computed by three-dimensional (3D) numerical simulation using commercial multiphysics software (COMSOL Multiphysics ver. 5.5, COMSOL AB, Sweden) and verified using an oxygen-sensing film (38). Simulations and measurements showed excellent agreement and yielded a sigmoidal oxygen gradient ranging from 0.4% to 21% along the medium channel *x*-direction when the gas mixtures containing 0% and 21% O_2_ were supplied to the left and right gas channels, respectively (red line in Fig. 3C). At the most hypoxic location at *x* = -1.6 mm, the slope of the oxygen concentration gradient inverted. Cells were harvested from the culture dishes and introduced into the medium channels in the microfluidic device at typically 10^6^ cells/ml (200 cells/mm^2^) with the PYG90 culture medium. After incubation for 20 min to adhere cells to the bottom surface of the medium channels, the PYG90 culture medium was replaced by fresh one. The device was mounted on the inverted microscope. Just after gas mixtures of oxygen and nitrogen were supplied at 30 ml/min to the left and right gas channels, time-lapse imaging was started.

### Time-lapse microscopy (bright field and fluorescence)

All time-lapse experiments were performed in an air-conditioned room with a constant temperature of 23°C on the microscope stage. Images of confined assays or inside microfluidic devices were collected at intervals compatible with cell displacement timescales (30 to 60 s/frame) for generally 2 hours using two distinct microscopes. A TE2000-E inverted microscope (Nikon) controlled with Micromanager (version 2.0 gamma) and equipped with a motorized stage, a 4X Plan Fluor objective lens (Nikon) and a Zyla camera (Andor) was used for most of the brightfield timelapse experiments of confinement and microfluidic assays. The same microscope was used to record the fluorescence signal of sensing films using a 4x Plan Fluor objective lens (Nikon), a X-Cite Series 120PC illumination lamp, a TRITC bandpass filter cube. A binocular MZ16 (Leica) controlled with LAS X software (version 3.4.2 Leica) and equipped with a TL3000 Ergo transmitted light base (Leica) operated in the one-sided darkfield illumination mode and a Leica LC/DMC camera (Leica) was used for long term, large field experiments (Fig. S2).

### Cell Tracking

For every assay (confinement and microfluidic assay), the individual cells were tracked using the open-source Python library *Trackpy* and standard scientific libraries *NumPy, pandas, matplotlib* and *SciPy* as described in (40). For each frame in the recorded time lapse series, we identified cell coordinates via the “Find Maxima” function in *ImageJ* (version 1.53q) (53) and then linked cell positions into trajectories using Trackpy across consecutive frames. Linking was performed with a conservative search range (10-40 pixels) depending on the frame rate and a short-term “memory” (4 frames across all datasets), allowing for the occasional dropout of cell centroids without prematurely ending trajectories. Occasionally, we segmented cells before the tracking using a machine-learning-based approach implemented in Ilastik (54).

### Analysis of cell displacements in microfluidic devices

For microfluidics experiments, we used cartesian coordinates. The *x* and *y*-axes were defined to be parallel and perpendicular to the medium channels, respectively. The origin was set at the center of the device. For each piece of trajectory *i*, the distance *dl*_*i*_*(t)* migrated during 1-min was computed as a function of the time *t*, after starting to flush gas in the gas channels. It was decomposed into the *x* and *y*-directional displacements *dl*_*x,i*_*(t)* and *dl*_*y,i*_*(t)*. We defined the average *x*-directional velocity 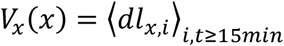 by averaging all cell displacements in Δ*x* = 100μ*m*wide slabs, all over *y* positions, as a function of *x* and over all times larger than 15 min which is the time necessary to establish a steady O_2_ gradient in this device (38).

### Density profiles and kymographs in confinement assays

For the rectangular stripe confinement assay, the origin *x=0* of the cartesian coordinates used was estimated from the stencil slit width and the initial border of the confined colony. Once the trajectories were tracked, the average *V*_*x*_(*x, t*) velocity was computed maps by averaging all 1-min *x*-directional cell displacements in Δ*x* = 20 µm wide slabs all over *y* positions as a function of *x*-position and time *t*. Densities maps *ρ*(*x, t*)were calculated by dividing by the slab surface the number of cells counted within Δ*x* = 20 µm wide slabs all over *y* positions. For the spot assay, the center of the colony was defined as the center of mass of all cells detected at all times. Cell positions were then turned into radial coordinates *r* and the average radial velocity *V*_*r*_(*r, t*) was computed by averaging all 1-min radial cell displacements in Δ*r* = 20 µm concentric crowns. Densities maps *ρ*(*r, t*)were calculated by dividing the cell number in each crown by the area of each crown. Both velocity and density maps were plotted as distance, time kymograph with a colorbar coding for the amplitude of the quantity plotted.

### Oxygen consumption measurements

Oxygen consumption was measured in sealed 6-mL glass bottles containing approximately 2 × 107 cells suspended in culture medium. Dissolved oxygen concentration was continuously monitored using a FireSting optical oxygen meter equipped with an OXROB3 probe (PyroScience, Germany). Cellular oxygen consumption was obtained from the initial linear decrease rate in oxygen concentration normalized by the number of cells.

### Mean-field ‘Go-or-Grow’ model

Numerical simulations were performed with the mean-field Go-or-Grow model previously developed for confined *Dictyostelium* colonies and implemented in Python (38, 39). The mean field model used for the aerotactic dispersion consisted of coupled reaction-diffusion equations (Eqs. 6 and 7) for both oxygen *C* and cell density *ρ*. The aerotactic flux was proportional to *χl*∇C/C, where both *χ*(*ρ*) and the oxygen-sensing cutoff function *f(C)* were determined experimentally (Fig. 4). The cellular diffusion coefficient was also fixed by the experimental *D(ρ)* (Fig. 4D). All remaining parameters, including oxygen consumption and proliferation rate, were independently measured experimentally (Fig. S5). Oxygen diffusion constant in PYG90 medium at 22°C was taken as 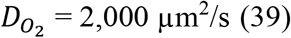. Boundary conditions reproduced the experimental geometry: *C*(*L*)=21% O_2_ with *L=*11 mm the position of the 22 mm coverslip border. Mesh sizes of Δ*x* = 10 µm and time interval Δ*t* = 0.05 min were used. To avoid negative O_2_ values a threshold *C*_1_ = 1% O_2_ was chosen below which cells consume oxygen relative to the local ambient oxygen concentrations. We also used a proliferation threshold below which cells stop dividing *C*_2_ = 0.2% O_2_ as recently measured (Carrère et al, 2022). This is also a slight modification to our initial Go-or-Grow model for which *C*_0_ = *C*_2_ (Cochet-Escartin et al, 2021).

## Supplementary figures and supplementary movies captions

**Fig. S1.**
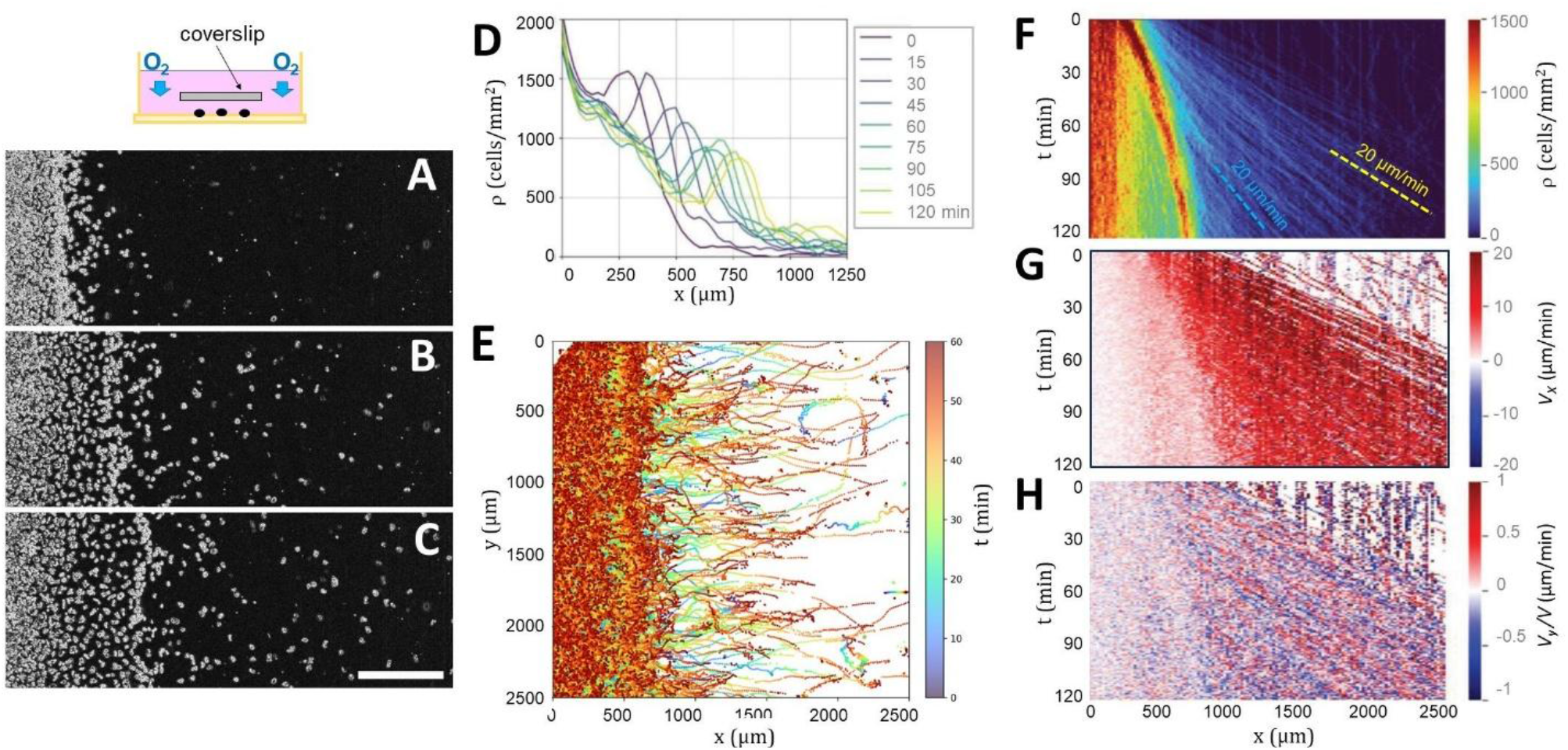
*Spreading dynamics of a confined rectangular stripe at high density of 1,200 cells/mm^2^*. (A–C) Snapshots at *t* = 0 (A), *t* = 60 min (B), and *t* = 120 min (C) showing the emergence of a diffuse front ring and ballistic leader cells expanding outward. Scale bar, 500 µm. (D) Cell-density profile averaged along the vertical y-direction as a function of the horizontal x-coordinate at various times. (E) Cell trajectories tracked for 60 min, color-coded by time. (F-H) Kymographs of cell density (F), velocity *V*_*x*_ along the horizontal x-coordinate (G) and normalized velocity *V*_*y*_ /*V* along the vertical coordinate where 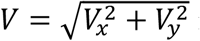 revealing the slow ring propagation with respect to ballistic cells and the strongly outwardly biased expansion dynamics while *V*_*y*_∼0. The yellow and blue dotted lines in (F) indicate ballistic cell trajectories with longitudinal velocities of 10 and 20 µm/min respectively.

**Fig. S2.**
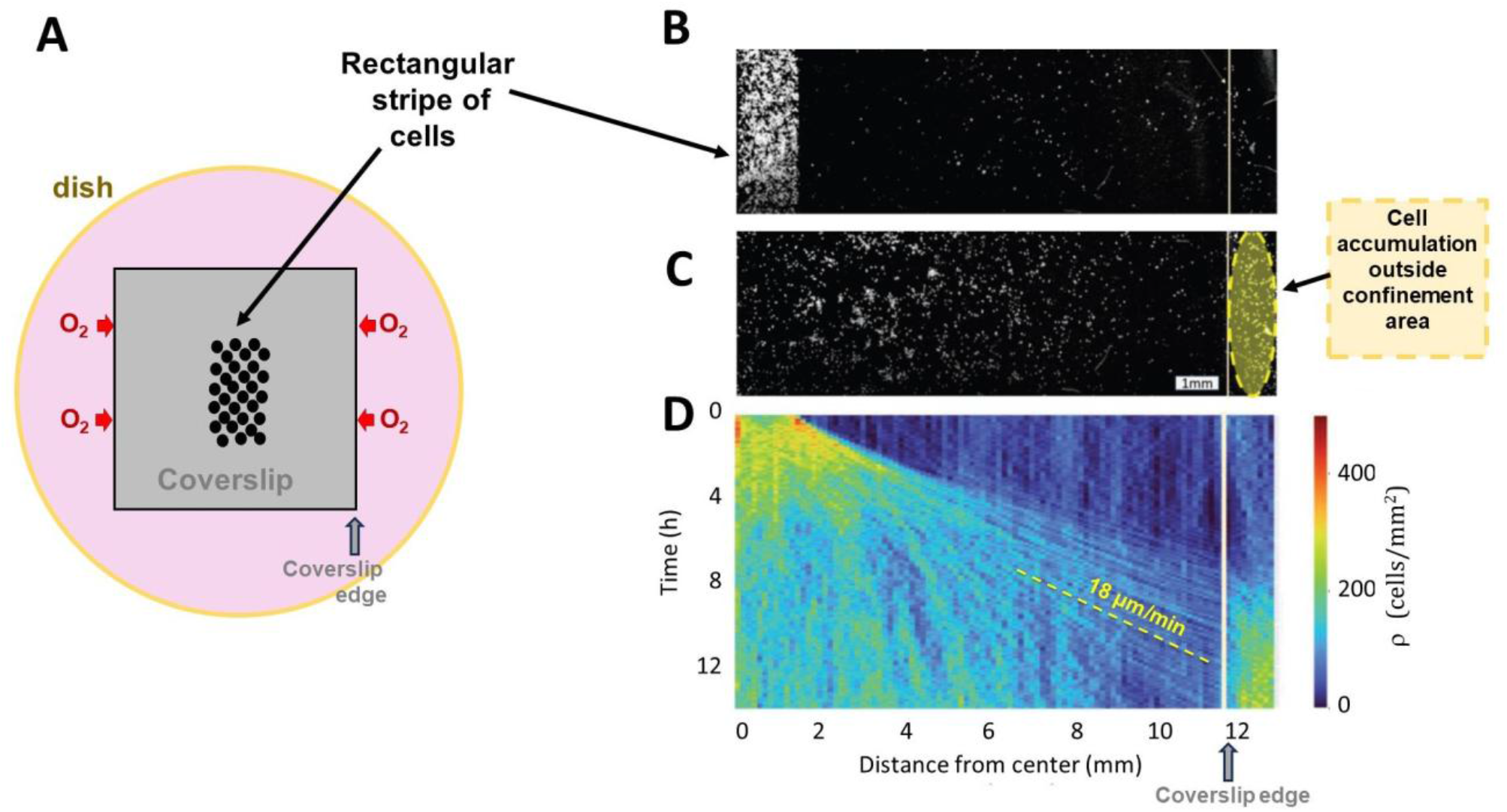
Long term outward escape of the confined area. (A) Schematic of the experiment: a rectangular cell stripe was confined beneath a 22 × 22 mm^2^ square coverslip, centered and oriented along the stripe, and imaged via wide-field microscopy to enable simultaneous observation of the initial stripe and the coverslip edge. (B) Initial low cell density stripe captured at *t*=0 immediately after confinement. The vertical white line indicates the coverslip edge, located approximately 11mm from the center of the rectangular stripe. (C) Final acquisition time at *t*=14 h. The yellow region highlights the accumulation of aerotactic cells escaping from beneath the coverslip. (D) Cell density kymograph with the vertical dotted line representing the position of the coverslip edge.

**Fig. S3.**
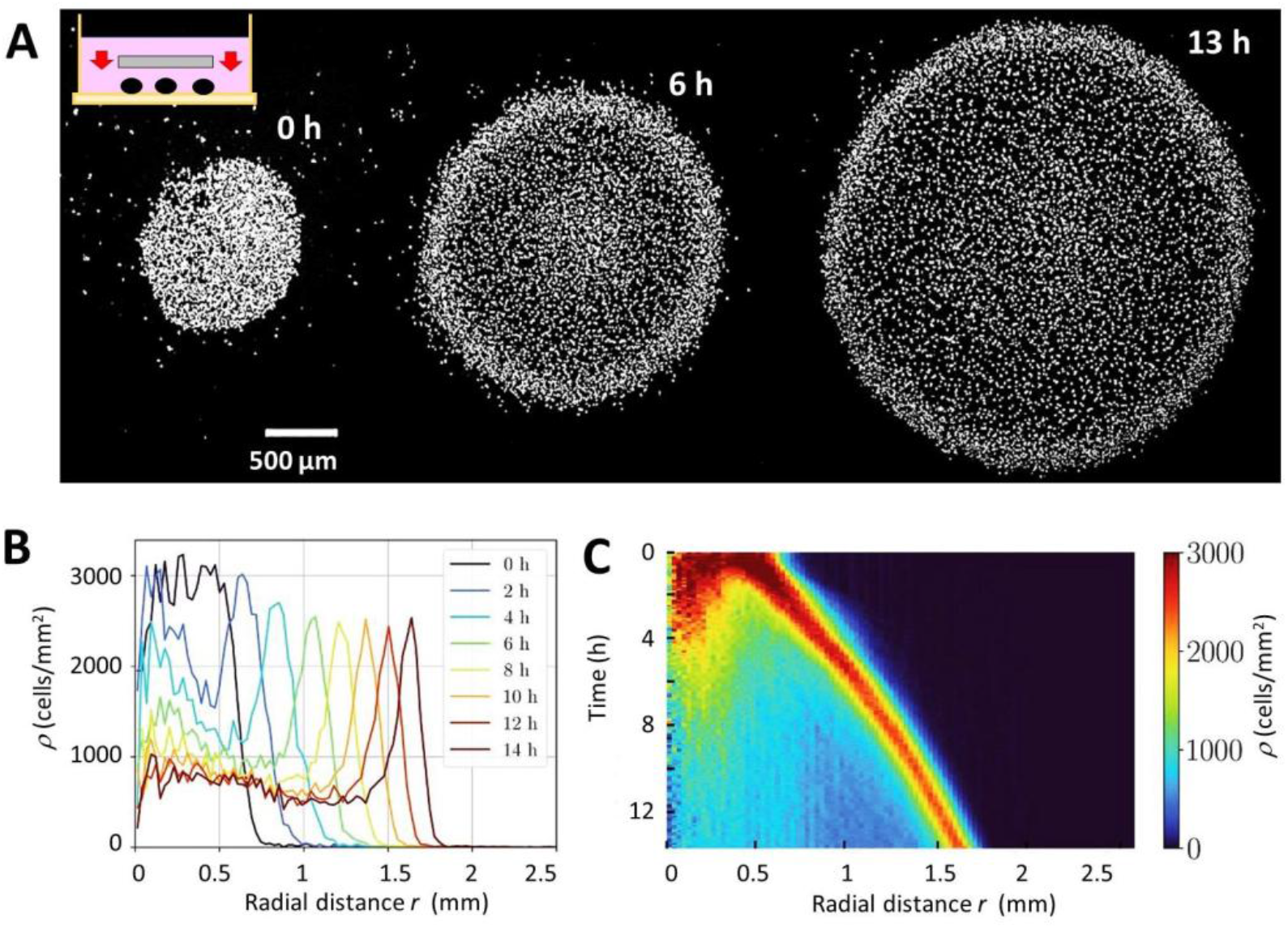
Collective spreading of *Dictyostelium discoideum* (Dd) in the confined spot assay. Approximately 3000 *Dd* cells were confined beneath a glass coverslip. (A) Time-lapse images showing the formation and long-term propagation of a dense, highly symmetric ring. (B) Radial cell-density profiles at successive times showing a nearly constant density peak propagating outward. (C) Radial density kymograph illustrating the steady outward migration of the ring at approximately 1 µm min^−1^ while maintaining a nearly constant peak density.

**Fig. S4.**
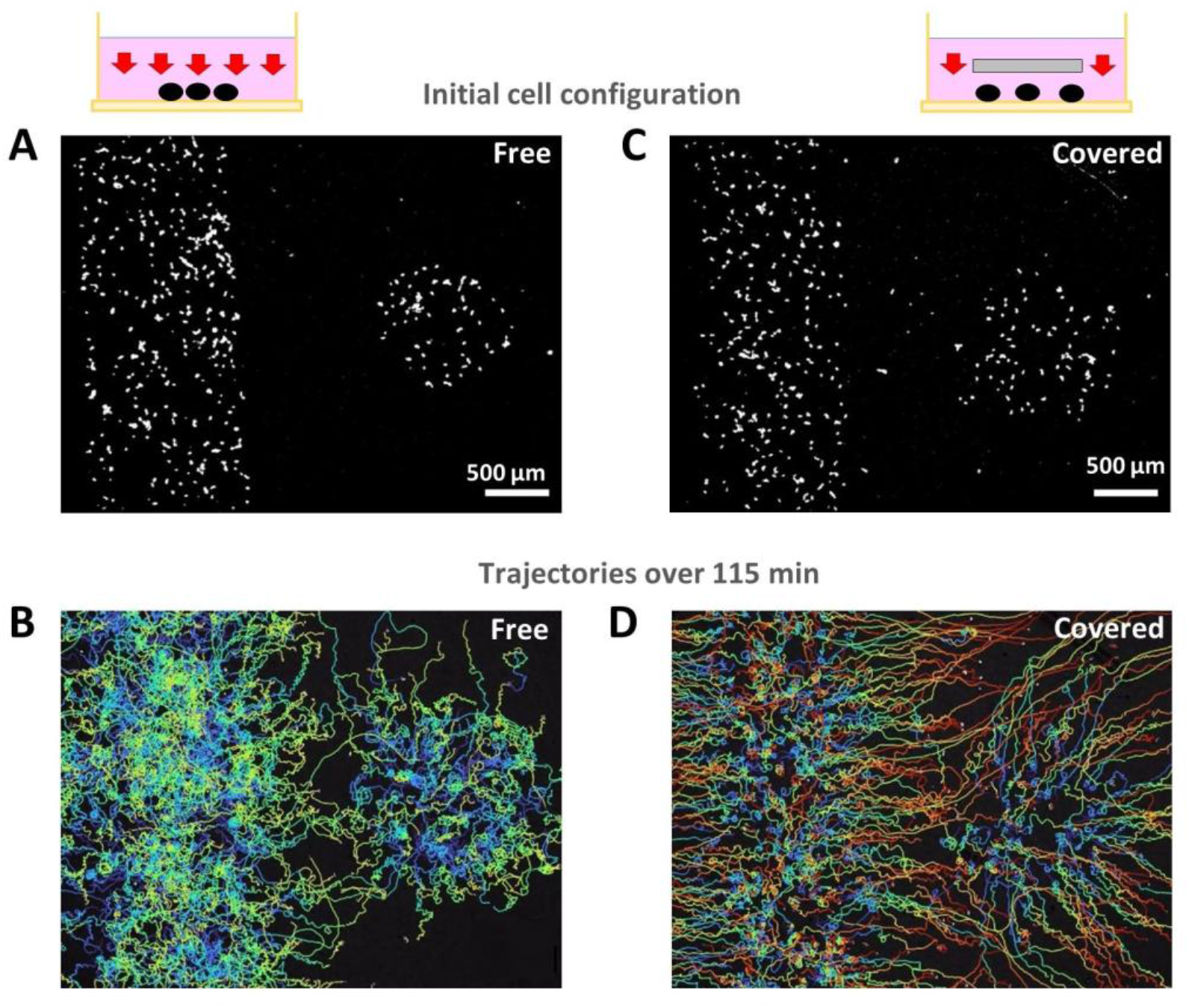
Confinement-induced long-range avoidance between opposing *Ac* colonies. (A,B) Unconfined (uncovered) control. A rectangular stripe and a circular spot of *Ac* were seeded at initial densities of *ρ* = 119 cells/mm^2^ (stripe) and *ρ* = 90 cells/mm^2^ (spot). (A) Initial snapshot at *t*=0. (B) Cell trajectories tracked over 115 min show diffusive spreading and free mixing upon contact, with no avoidance beyond local collision-induced reorientations. (C,D) Confined (covered) condition. The same geometry was reproduced beneath a coverslip, with initial densities of *ρ* = 118 cells/mm^2^ (stripe) and *ρ* = 87 cells/mm^2^ (spot). (C) Initial snapshot at *t*=0. (D) Cell trajectories tracked over 115 min reveal pronounced avoidance: stripe cells migrate primarily along the ±x, spot cells expand radially outward, and both populations bend their trajectories away from the opposing colony upon approach. Confinement thus triggers a long-range guided avoidance response, consistent with an aerotactic mechanism.

**Fig. S5.**
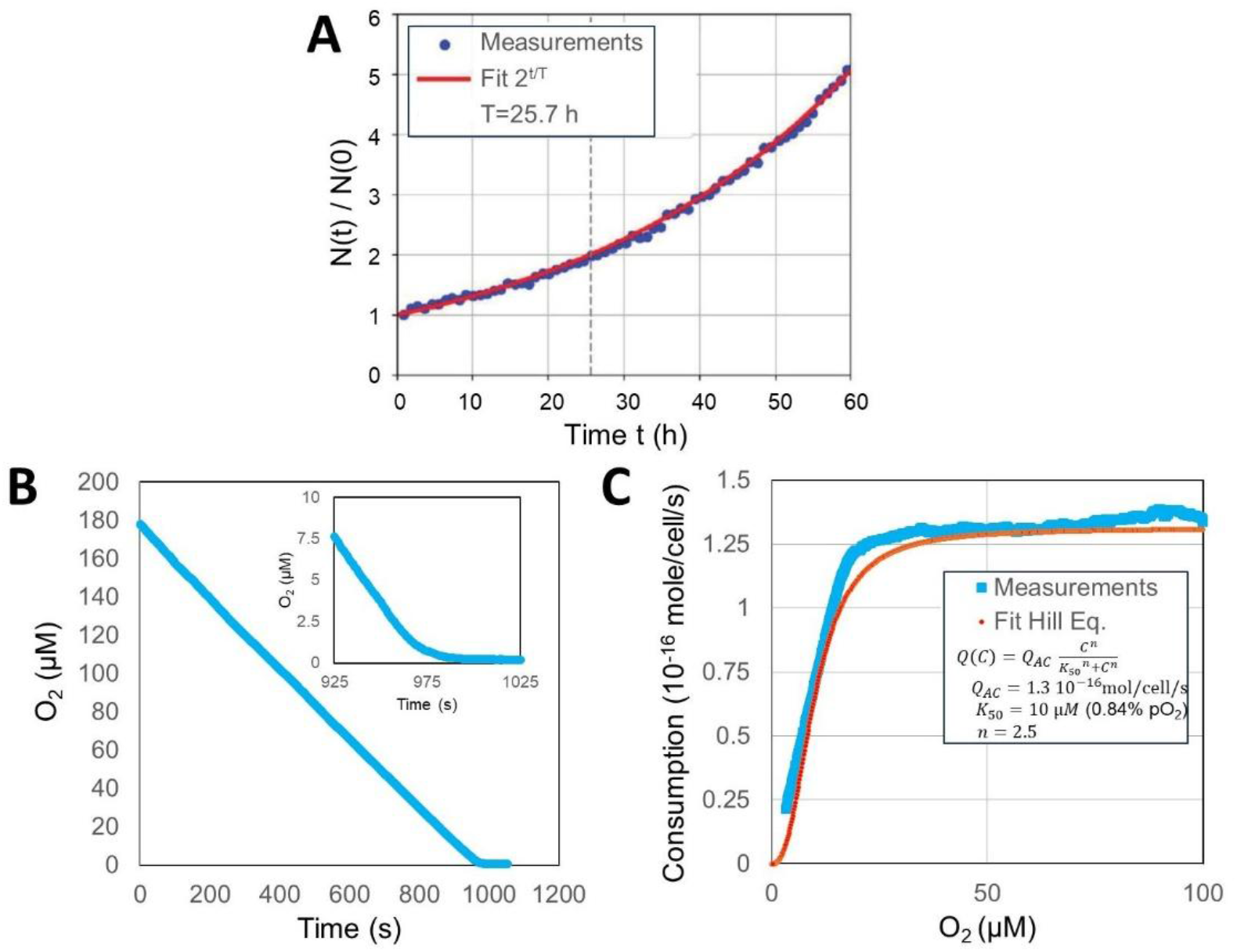
*Ac* other experimental parameters needed for the models. (A) Cell proliferation experiments (*i*.*e*., number of cells *N(t)* at time t divided by initial number *N(0)*) presenting a nearly 24 h doubling time (vertical dotted line). (B) Oxygen drop in a closed bottle filled with *Ac* cells and (C) corresponding calculated oxygen consumption fitted with the law written in the graph. The consumption is fitted with Hill equation giving a maximum consumption of 1.3 10^-16^ mole/cell/s, a concentration at half-maximum of 10 µM and a Hill coefficient of 2.5 (equivalent to 0.84 % pO_2_) (Lavrinenko. *et al*. Mathematical models describing oxygen binding by hemoglobin. *Biophys Rev* **15**, 1269–1278 (2023))

**Fig. S6.**
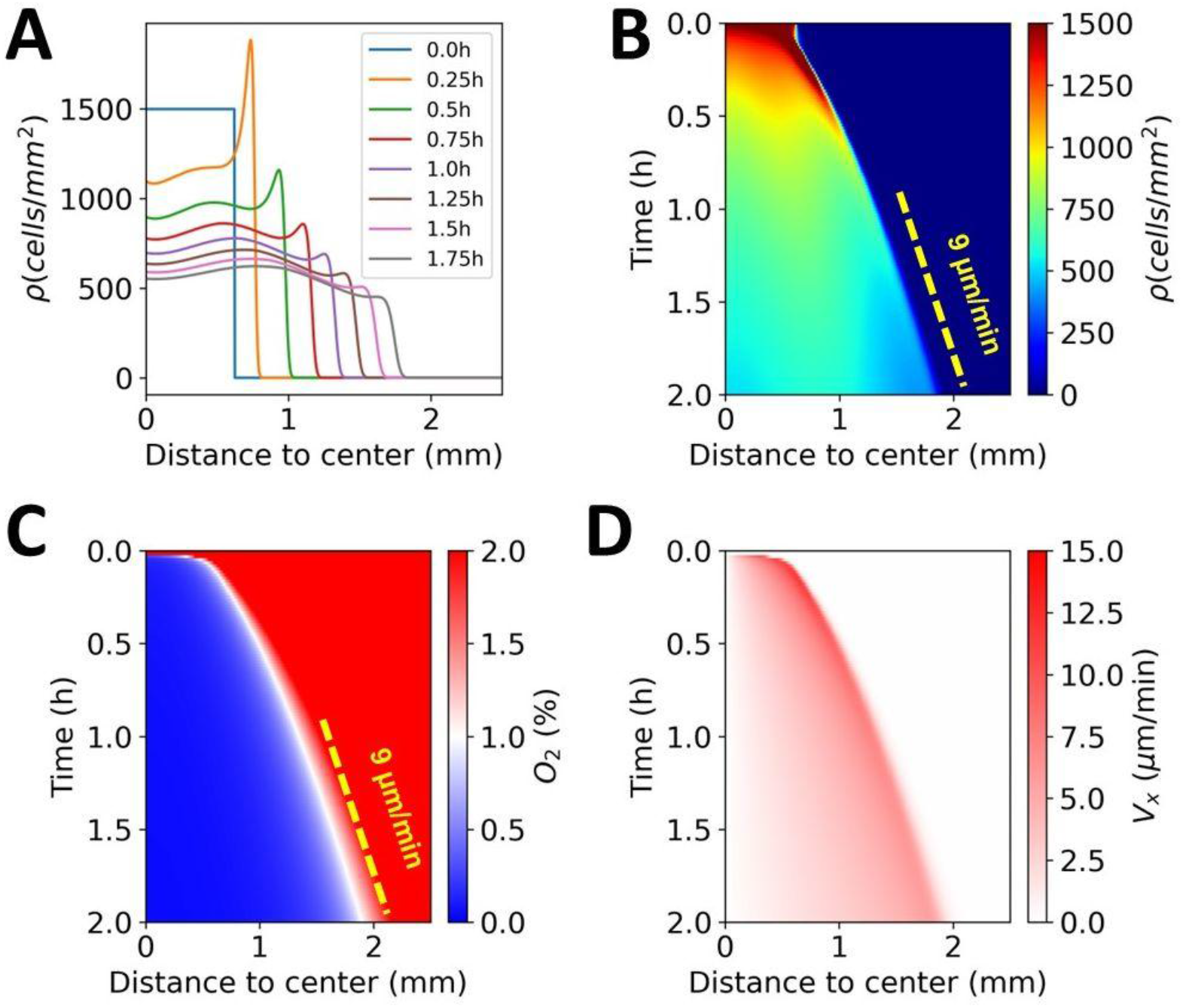
**Modelling Ac spreading from a covered rectangular stripe** using a small constant cellular diffusion constant (*D*_*0*_=72 µm^2^/min) and aerotactic bias speed *X*_*0*_=37 µm/min.

**Movie S1. Spreading dynamics of an** *Acanthamoeba castellanii* **confined circular spot generated using a 1.25mm diameter stencil**. The spot was formed using a 1.25mm diameter punched stencil, seeded at a cell density *ρ* = 1200 cells/mm^2^ and covered with a coverglass. The movie highlights the four spreading zones: crowded core, intermediate region, diffuse ring, and ballistic cells. The full field of view is 4mm x 3.2mm.

**Movie S2**. *Spreading dynamics of a covered rectangular stripe of cells. The stripe was formed using a rectangular stencil at an initial density of 1,200 cells mm*^*−2*^. The movie show the emergence of a diffusive front ring and ballistic leader cells expanding outwardly.

**Movie S3. Long term** *spreading a covered rectangular stripe of cells*. A rectangular cell stripe was confined within a square coverslip measuring 22 mm in width, centered and oriented along the stripe and observed with a wide-field microscopy allowing simultaneous observation the initial stripe and the coverglass border.

**Movie S4. Confinement-induced avoidance between opposing *Ac* colonies**. A rectangular stripe and a circular spot of *Ac* were spotted at initial densities of *ρ* = 119 cells mm^−2^ (stripe) and *ρ* = 90 cells mm^−2^ (spot), covered by a coverglass (right side) or not (free case, left side) and observed by time lapse during 2 hours. The full field of view is 4mm x 3.2mm.

**Movie S5. Coupled cell-density and oxygen dynamics reveal aerotaxis-driven spreading in confined *Ac* stripes**. (Top left) Time-lapse of a confined rectangular stripe of *Ac* (initial density *ρ* = 1200 cells mm^−2^), showing outward propagation of a diffuse ring and fast, outward-oriented ballistic cells. The stripe is covered by a glass coverslip coated with a ∼50-μm-thick O_2_-sensing film (PDMS embedding the porphyrin dye PtTFPP). (Bottom left) Reconstructed O_2_ map beneath the coverslip at t = 90 min measured with from porphyrin phosphorescence and (right) corresponding density (red) and oxygen (red) rofile along the horizontal direction x.

**Movie S6. Interaction of outsider cells with leader cells under confined conditions**. (Left) trajectories (right) local O_2_ map measured with sensing films (same experiment as Fig. 2 and Movie M5) in the region of outsider cells initially located away from the initial cellular stripe. As leader cells approach from the left (−x direction), local O_2_ levels decrease, and the outsider cell begins directed migration toward the leader pack (+x direction).

## Supplementary Information Appendix (SI Appendix)

## Emergent transport coefficients quantitatively link microscopic collisions to collective aerotactic migration

### From collision-induced reorientations to density-dependent aerotaxis

To provide a microscopic rationale for the density dependence of the aerotactic coefficient *χ*(*ρ*), we consider a minimal one-dimensional persistent random-walk model. This construction provides a possible microscopic interpretation of the functional dependence used in the macroscopic model, by showing how it can arise naturally when cell-cell collisions modulate directional persistence.

Cells move with constant speed *V*_0_ either in the positive or negative direction. We denote these two states by + and −, respectively. As established in our previous work (35), we account for the effect of cell-cell collisions at a mean-field level by introducing a density-dependent reorientation rate,

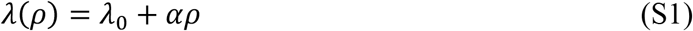

where *λ*_0_ is the intrinsic reorientation rate, observed at very low densities, and *αρ* represents the collision-induced contribution to the reorientation rate, with *ρ* denoting the local cell density and *α* a coefficient. In (35), a geometric interpretation was obtained from mean-free path theory, and we obtained the formula 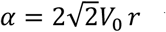, where *r* represents the radius of the cell.

Within this minimal description, we allow here aerotaxis to act through two complementary mechanisms. First, the aerotactic signal may modulate the reorientation frequency, thereby changing the persistence time depending on the direction of motion: we refer to this as *temporal bias*. Such a mechanism is analogous to temporal sensing in bacterial chemotaxis, where reorientation frequencies are modulated according to changes in the sensed chemical environment along the cell trajectory, thereby prolonging runs in favorable directions and shortening runs in unfavorable directions (16, 30). Second, the direction selected after a reorientation event may itself be biased toward the favorable direction: we refer to this as *spatial bias*. Such a mechanism is analogous to spatial gradient sensing in eukaryotic chemotaxis, where spatial differences in the sensed chemoattractant bias cell polarization and migration toward higher concentrations (15). Together, these two mechanisms provide a simple microscopic explanation for aerotaxis. The dependence on cell density then arises through the density-dependent reorientation rate *λ*(*ρ*).

We denote the local dimensionless aerotactic signal by

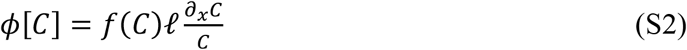

where *C* is the oxygen concentration, *f*(*C*) is an oxygen-dependent cutoff function, and *l* is a characteristic length introduced to adimensionalize the logarithmic oxygen gradient. The quantity *ϕ*[*C*] represents the local strength and direction of the aerotactic cue combining the logarithmic oxygen gradient with the oxygen-dependent cutoff. We use square brackets to emphasize that, in general, *ϕ*[*C*] may depend on the entire oxygen field *C* rather than only on its value at a single point. In the present model, however, this dependence is local and involves only the values of *C* and its spatial derivative ∂_*x*_*C*. The sign of *ϕ*[*C*] determines the favorable direction of migration.

### Temporal bias

As a first possible contribution to aerotaxis, we consider a modulation of the reorientation frequency by the direction of migration. Cells moving in the favorable direction are assigned to a reorientation rate *λ*_+_ smaller than the reorientation rate *λ*_−_ of cells moving in the unfavorable direction. We write

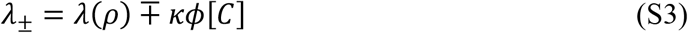

where *κ* > 0 measures the strength of the temporal bias. Thus, when *ϕ*[*C*] > 0runs in the positive direction are prolonged whereas runs in the negative direction are shortened, generating a net bias of the cell displacement towards increasing oxygen concentration.

### Spatial bias

As a second possible contribution to aerotaxis, we consider a bias in the direction selected after a reorientation event. Let *p*_+_ and *p*_−_ denote the probabilities that, following a reorientation, the cell chooses the positive or negative direction, respectively We write

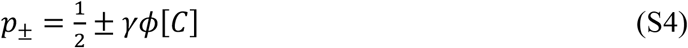

where *γ* > 0 measures the strength of the spatial bias. Thus, when *ϕ*[*C*] > 0

The two mechanisms act in complementary ways: the spatial bias modifies the direction selected after reorientation, whereas the temporal bias modifies the time spent moving in each direction.

### Stationary directional distribution and mean drift

A cell moving in the positive (resp. negative) direction switches to the negative (resp. positive) direction only when a reorientation event occurs, and the negative (resp. positive) direction is subsequently selected. The corresponding effective switching rate from positive to negative (resp. negative to positive) is therefore

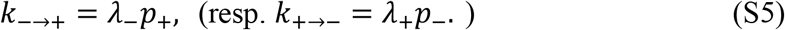

The stationary fractions of time moving in the two directions are

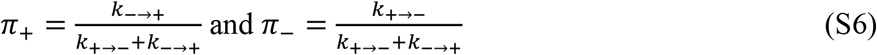

The resulting mean aerotactic velocity is therefore

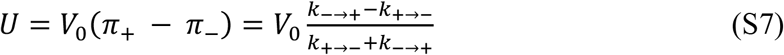

Substituting the expressions above, together with the definitions of *λ*_±_ and *p*_±_, gives the exact stationary drift

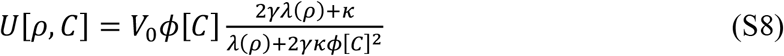

Introducing the dimensionless correction

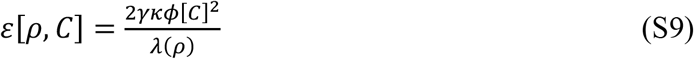

we may rewrite the exact stationary drift as

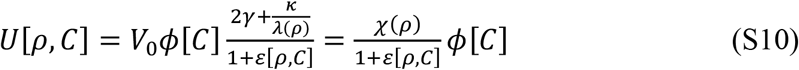

Where

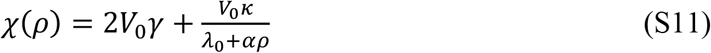

This expression provides a possible microscopic interpretation of the density dependence used in the macroscopic model. In the regime where *ε* ≪ 1, the correction from the denominator remains small and the aerotactic velocity is approximated by

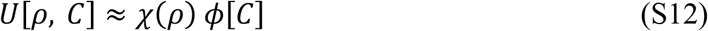

Within this approximation, the spatial and temporal biases contribute additively to the effective aerotactic coefficient, with a density-independent spatial contribution and a temporal contribution proportional to 1/*λ*(*ρ*).

A remarkably similar density dependence of the chemotactic coefficient was introduced phenomenologically for dense populations of E. coli in Phan *et al*. (27). Rewriting their chemotactic drift in our notation, it takes the form

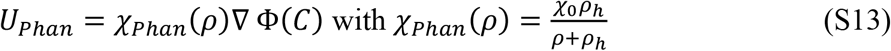

where Φ(*C*) (which is distinct from the previous *ϕ*[*C*]) describes the chemical sensing response and *ρ*_*h*_ is the characteristic cell density at which the chemotactic coefficient is reduced to one half of its low-density value. Thus, *χ*_*Phan*_(*ρ*) exhibits the same reciprocal-linear dependence on cell density as the temporal contribution 1/*λ*(*ρ*) in the present model. Related experiments and agent-based simulations by Colin *et al*. (47) associated the reduction of chemotactic efficiency at high cell density with cell reorientations induced predominantly by hydrodynamic interactions, which shorten directional persistence and thereby weaken the efficiency of temporal chemotactic sensing. In contrast to the present model, this description does not include an additional density-independent spatial bias contribution, consistent with the predominantly temporal sensing strategy of *E. coli* (16, 30).

### Joint parametrization of diffusion and aerotactic drift

Motivated by the expression above, we view the collision-induced increase in reorientation frequency as a possible microscopic scenario leading to the observed density dependence and use it to motivate the phenomenological form adopted in the macroscopic model. We therefore assume in the main text that the aerotactic coefficient takes the form

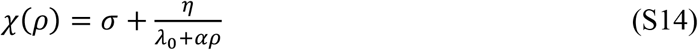

where *σ* = 2 *V*_0_*γ, η* = *V*_0_ *κ*. Importantly, the same density-dependent reorientation rate *λ*(*ρ*) = *λ*_0_ + *αρ* also determines the undirected diffusive random motility. From the mean-free-path argument introduced in our previous work (35), the diffusion coefficient in two spatial dimensions is

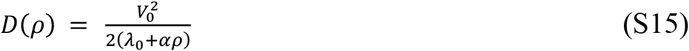

We therefore fit the diffusion coefficient *D*(*ρ*) and aerotactic coefficient *χ*(*ρ*) simultaneously from the experimental data, using the same density-dependent reorientation rate *λ*_0_ + *αρ* in both coefficients. The resulting parameter values are reported in the main text.

We can also assess a posteriori the consistency of the approximation in Eq. (S12). Rewriting Eq. (S9) using Eq. (S2) and *σ* = 2 *V*_0_*γ, η* = *V*_0_ *κ*, we find

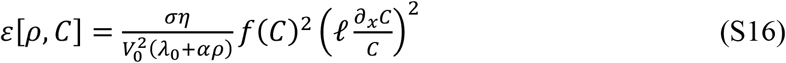

This provides a direct measure of the consistency of the approximation of Eq. (S12): once the common transport parameters have been estimated from the measurements of *D*(*ρ*) and *χ*(*ρ*), the magnitude of *ε* can be evaluated over the experimentally relevant ranges of cell density and oxygen gradients.

### Generalization to a continuous set of directions

The two-velocity construction above provides an exact and tractable expression for the stationary drift, including a correction controlled by *ε*[*ρ, C*]. We now briefly generalize the argument to a continuous set of velocities. The purpose of this calculation is not to reproduce the specific correction *ε*[*ρ, C*] obtained in the two-velocity model, but rather to verify that the leading-order dependence *χ*(*ρ*) = *σ* + *η*/*λ*(*ρ*) is not an artifact of restricting cell motion to two directions.

We consider cells moving in two spatial dimensions with constant speed *V*_0_ and direction 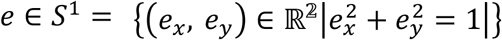We introduce the vector-valued aerotactic signal

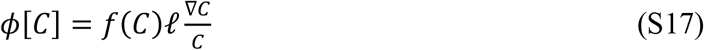

As in the two-velocity model, we allow aerotaxis to act through both the reorientation frequency and the direction selected after a reorientation event. We write the direction-dependent reorientation frequency as

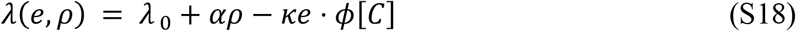

while the direction after reorientation is sampled from a probability distribution *M*_*ϕ*_(*e*) on *S*^1^. We assume that in the absence of an aerotactic signal this distribution is isotropic, and that its mean direction responds linearly to the signal,

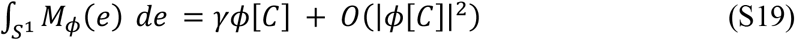

The precise form of *M*_*ϕ*_(*e*) is otherwise unspecified.

For fixed local values of *ρ* and *C*, a direction *e* is selected with probability density *M*_*ϕ*_(*e*) and is retained for an average time 1/*λ*(*e, ρ*). The stationary distribution of directions is therefore proportional to the product of these two quantities,

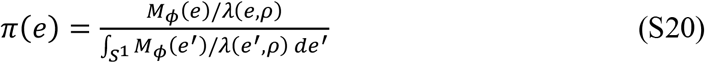

The corresponding mean migration velocity is

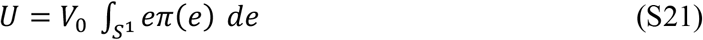

To compare this expression with the reduced two-velocity law, we expand only to first order in the aerotactic signal. From the definition of *λ*(*e, ρ*),

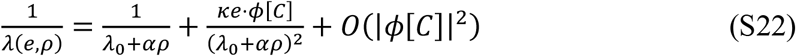

Therefore

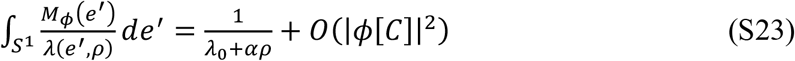

And

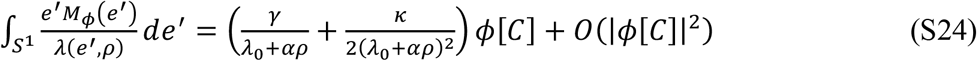

Using those two expressions with Eq. (S21) and Eq. (S20) then leads to

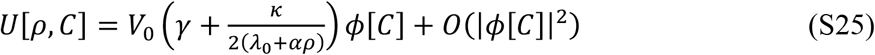

Thus, the same leading-order structure as in the two-velocity model is recovered:

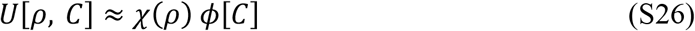

Within this first-order description, the spatial and temporal contributions are again additive: the spatial contribution is independent of the baseline reorientation frequency, whereas the temporal contribution scales as 1/(*λ*_0_ + *α ρ*). The expression of *χ*(*ρ*) in Eq. (S25) differs by a factor 1/2 from the two-velocity model in Eq. (S11). This is due to an averaging over directions on *S*^1^, but such geometrical factors can be absorbed into the effective parameters *σ* and *η*.

Because the expansion above is restricted to first order in *ϕ*[*C*], higher-order corrections in the aerotactic signal, such as the quantity *ε*[*ρ, C*] obtained in the exact two-velocity model do not appear at this order. Their detailed form depends on the specific angular reorientation distribution *M*_*ϕ*_. The role of the continuous-velocity calculation is therefore to show that the leading-order functional dependence *χ*(*ρ*) = *σ* + *η*/(*λ*_0_ + *αρ*) persists beyond the two-velocity model.

## References

1. P. Friedl, D. Gilmour, Collective cell migration in morphogenesis, regeneration and cancer. Nat. Rev. Mol. Cell Biol. 10, 445–457 (2009).

2. A. D. Luster, R. Alon, U. H. von Andrian, Immune cell migration in inflammation: present and future therapeutic targets. Nat. Immunol. 6, 1182–1190 (2005).

3. P. Martin, Wound Healing--Aiming for Perfect Skin Regeneration. Science 276, 75–81 (1997).

4. E. S. Gloag, et al., Self-organization of bacterial biofilms is facilitated by extracellular DNA. Proc. Natl. Acad. Sci. 110, 11541–11546 (2013).

5. F. W. Rossine, G. T. Vercelli, C. E. Tarnita, T. Gregor, Structured foraging of soil predators unveils functional responses to bacterial defenses. Proc. Natl. Acad. Sci. U. S. A. 119, e2210995119 (2022).

6. C. Carmona-Fontaine, et al., Contact inhibition of locomotion in vivo controls neural crest directional migration. Nature 456, 957–961 (2008).

7. J. D’alessandro, et al., Contact enhancement of locomotion in spreading cell colonies. Nat. Phys. 13, 999–1005 (2017).

8. T. Fujimori, A. Nakajima, N. Shimada, S. Sawai, Tissue self-organization based on collective cell migration by contact activation of locomotion and chemotaxis. Proc. Natl. Acad. Sci. 116, 4291–4296 (2019).

9. Y. Li, et al., In Vivo Quantitative Imaging Provides Insights into Trunk Neural Crest Migration. Cell Rep. 26, 1489–1500.e3 (2019).

10. T. Hiraiwa, Dynamic Self-Organization of Idealized Migrating Cells by Contact Communication. Phys. Rev. Lett. 125, 268104 (2020).

11. W. Alt, Biased random walk models for chemotaxis and related diffusion approximations. J. Math. Biol. 9, 147–177 (1980).

12. T. Hillen, K. J. Painter, A user’s guide to PDE models for chemotaxis. J. Math. Biol. 58, 183–217 (2009).

13. H. G. Othmer, S. R. Dunbar, W. Alt, Models of dispersal in biological systems. J. Math. Biol. 26, 263–298 (1988).

14. M. J. Schnitzer, Theory of continuum random walks and application to chemotaxis. Phys. Rev. E 48, 2553–2568 (1993).

15. T. Jin, Gradient sensing during chemotaxis. Curr. Opin. Cell Biol. 25, 532–537 (2013).

16. H. C. Berg, D. A. Brown, Chemotaxis in Escherichia coli analysed by Three-dimensional Tracking. Nature 239, 500–504 (1972).

17. C. A. Parent, P. N. Devreotes, A Cell’s Sense of Direction. Science 284, 765–770 (1999).

18. K. F. Swaney, C.-H. Huang, P. N. Devreotes, Eukaryotic Chemotaxis: A Network of Signaling Pathways Controls Motility, Directional Sensing, and Polarity. Annu. Rev. Biophys. 39, 265–289 (2010).

19. J. H. R. Wheeler, K. R. Foster, W. M. Durham, Individual bacterial cells can use spatial sensing of chemical gradients to direct chemotaxis on surfaces. Nat. Microbiol. 9, 2308–2322 (2024).

20. Bonner J.T., The Cellular Slime Molds, P. U. Press, Ed. (1967).

21. R. H. Kessin, Dictyostelium: Evolution, Cell Biology, and the Development of Multicellularity, C. U. Press, Ed. (2001).

22. R. H. Insall, Receptors, enzymes and self-attraction as autocrine generators and amplifiers of chemotaxis and cell steering. Curr. Opin. Cell Biol. 81, 102169 (2023).

23. J. Adler, Chemotaxis in Bacteria. Science 153, 708 (1966).

24. L. Tweedy, et al., Seeing around corners: Cells solve mazes and respond at a distance using attractant breakdown. Science 369 (2020).

25. A. J. Muinonen-Martin, et al., Melanoma Cells Break Down LPA to Establish Local Gradients That Drive Chemotactic Dispersal. PLoS Biol. 12 (2014).

26. A. Kiran, et al., Macrophages self-generate and refine chemotactic gradients during migration towards complement C5a. PLOS Biol. 24, e3003728 (2026).

27. E. F. Keller, L. A. Segel, Initiation of Slime Mold Aggregation Viewed as an Instability. J. Theor. Biol. 26, 399–415 (1970).

28. E. F. Keller, L. A. Segel, Traveling bands of chemotactic bacteria: A theoretical analysis. J. Theor. Biol. 30, 235–248 (1971).

29. C. S. Patlak, Random walk with persistence and external bias. Bull. Math. Biophys. 15, 311–338 (1953).

30. J. Saragosti, et al., Directional persistence of chemotactic bacteria in a traveling concentration wave. Proc. Natl. Acad. Sci. 108, 16235–16240 (2011).

31. L. Tweedy, D. A. Knecht, G. M. Mackay, R. H. Insall, Self-Generated Chemoattractant Gradients: Attractant Depletion Extends the Range and Robustness of Chemotaxis. PLoS Biol. 14, e1002404 (2016).

32. T. V. Phan, et al., Direct measurement of dynamic attractant gradients reveals breakdown of the Patlak–Keller–Segel chemotaxis model. Proc. Natl. Acad. Sci. 121, e2309251121 (2024).

33. H. F. Bunn, R. O. Poyton, Oxygen sensing and molecular adaptation to hypoxia. Physiol. Rev. 76, 839–885 (1996).

34. T. W. Engelmann, Neue methode zur untersuchung der sauerstoffausscheidung pflanzlicher und tierischer organismen (New method for investigation of oxygen-searching plant and animal organisms). Pflugers Arch Gesammte Physiol 25, 285–292 (1881).

35. K. M. Winn, D. G. Bourne, J. G. Mitchell, Vibrio coralliilyticus Search Patterns across an Oxygen Gradient. PLOS ONE 8, 1–8 (2013).

36. M. Deygas, et al., Redox regulation of EGFR steers migration of hypoxic mammary cells towards oxygen. Nat. Commun. 9:4545 (2018).

37. I. Mikaelian, et al., EGFR-dependent aerotaxis is a common trait of breast tumour cells. J. Exp. Clin. Cancer Res. 41, 324 (2022).

38. O. Cochet-Escartin, et al., Hypoxia triggers collective aerotactic migration in Dictyostelium discoideum. eLife 10, e64731 (2021).

39. S. Hirose, et al., The aerotaxis of Dictyostelium discoideum is independent of mitochondria, nitric oxide and oxidative stress. Front. Cell Dev. Biol. Volume 11-2023 (2023).

40. N. Ghazi, et al., Density-dependent cell migration in the absence of social interactions: a case study of Acanthamoeba castellanii. Eur. Phys. J. E 48, 42 (2025).

41. R. Siddiqui, R. Dudley, N. A. Khan, Acanthamoeba differentiation: a two-faced drama of Dr Jekyll and Mr Hyde. Parasitology 139, 826–834 (2012).

42. N. A. Kuburich, N. Adhikari, J. A. Hadwiger, Acanthamoeba and Dictyostelium Use Different Foraging Strategies. Protist 167, 511–525 (2016).

43. F. L. Schuster, M. Levandowsky, Chemosensory Responses of Acanthamoeba castellanii: Visual Analysis of Random Movement and Responses to Chemical Signals. J. Eukaryot. Microbiol. 43, 150–158 (1996).

44. H. C. Berg, Ed., “Cell Populations” in E. Coli in Motion, (Springer New York, 2004), pp. 19–30.

45. M. J. Simpson, et al., Quantifying the roles of cell motility and cell proliferation in a circular barrier assay. J. R. Soc. Interface 10, 20130007 (2013).

46. S. Hirose, J. Rieu, O. Cochet-escartin, C. Anjard, K. Funamoto, The Oxygen Gradient in Hypoxic Conditions Enhances and Guides Dictyostelium discoideum Migration. Processes 10, 318 (2022).

47. R. Colin, K. Drescher, V. Sourjik, Chemotactic behaviour of Escherichia coli at high cell density. Nat. Commun. 10, 5329 (2019).

48. M. C. Marchetti, et al., Hydrodynamics of soft active matter. Rev. Mod. Phys. 85, 1143–1189 (2013).

49. F. Gu, B. Guiselin, N. Bain, I. Zuriguel, D. Bartolo, Emergence of collective oscillations in massive human crowds. Nature 638, 112–119 (2025).

50. C. Bechinger, et al., Active particles in complex and crowded environments. Rev Mod Phys 88, 45006 (2016).

51. R. J. Neff, Purification, Axenic Cultivation, and Description of a Soil Amoeba, Acanthamoeba sp. J. Protozool. 4, 176–182 (1957).

52. S. Hirose, J.-P. Rieu, O. Cochet-Escartin, C. Anjard, K. Funamoto, The Oxygen Gradient in Hypoxic Conditions Enhances and Guides Dictyostelium discoideum Migration. Processes 10 (2022).

53. J. Schindelin, et al., Fiji: an open-source platform for biological-image analysis. Nat. Methods 9, 676–682 (2012).

54. S. Berg, et al., ilastik: interactive machine learning for (bio)image analysis. Nat. Methods 16, 1226–1232 (2019).

